# Piezo1 Induces Local Curvature in a Mammalian Membrane and Forms Specific Protein-Lipid Interactions

**DOI:** 10.1101/787531

**Authors:** Amanda Buyan, Charles D. Cox, James Rae, Jonathan Barnoud, Jinyuan Li, Jasmina Cvetovska, Michele Bastiani, Hannah S.M. Chan, Mark P. Hodson, Boris Martinac, Robert G Parton, Siewert J. Marrink, Ben Corry

**Affiliations:** Research School of Biology, Australian National University, Acton, ACT 2601, Australia; Victor Chang Cardiac Research Institute, Lowy Packer Building, 405 Liverpool St., Darlinghurst, NSW 2010, Australia; St Vincent’s Clinical School, University of New South Wales, Darlinghurst, NSW 2010, Australia; Groningen Biomolecular Sciences and Biotechnology Institute, University of Groningen, Groningen, The Netherlands; Institute for Molecular Bioscience, University of Queensland, QLD 4072, Australia; Freedman Foundation Metabolomics Facility, Victor Chang Innovation Centre, Victor Chang Cardiac Research Institute, Lowy Packer Building, 405 Liverpool Street, Darlinghurst NSW 2010, Australia; School of Pharmacy, University of Queensland, QLD 4102, Australia; Centre for Microscopy and Microanalysis, University of Queensland, QLD 4072, Australia

**Keywords:** mechanosensation, mechanically-gated channels, electrophysiology, electron microscopy, molecular dynamics simulation, protein-lipid interaction

## Abstract

Touch, hearing, and blood pressure control require mechanically-gated ion channels that convert mechanical stimuli into electrical currents. Piezo1 and Piezo2 were recently identified as essential eukaryotic mechanically-gated ion channels, yet how they respond to physical forces remains poorly understood. Here we use a multi-disciplinary approach to interrogate the interaction of Piezo1 with its lipid environment. We show that individual Piezo1 channels induce significant local curvature in the membrane that is magnified in a cooperative manner to generate larger curved ‘Piezo1 pits.’ Curvature decreases under lateral membrane tension, consistent with a hypothesis that force detection can involve sensing changes to local curvature. The protein alters its local membrane composition, enriching specific lipids and forming essential binding sites for phosphoinositides and cholesterol that are functionally relevant and often related to Piezo1-mediated pathologies. Finally, we show that Piezo1 alters the expression of lipid-regulating proteins and modifies the cellular lipidome. In short, we find that lipids influence Piezo1 activity and Piezo1 influences the local morphology and composition of the bilayer as well as the cellular lipidome.

## Introduction

Piezo1 is a long-sought after molecular force sensor in eukaryotes (Coste et al., 2010). This non-selective cation channel is ubiquitously expressed, and is responsible for primary mechanosensitive currents in a myriad of cells and tissues (Servin-Vences et al., 2017; Zeng et al., 2018). Its presence in mammals is essential, exemplified by the embryonic lethality of its global deletion in mice (Li et al., 2014; Ranade et al., 2014). Moreover, Piezo1 gene variants have been linked to a number of human pathologies caused by either gain-of-function (e.g. hereditary xerocytosis (Albuisson et al., 2013; Zarychanski et al., 2012)) or loss-of-function (e.g. generalized lymphatic dysplasia (Lukacs et al., 2015)) mutations. In all likelihood, this list of known pathologies will grow given the broad expression pattern of Piezo1. The gain-of-function Piezo1 phenotype, identified by its impact on red blood cell (RBC) morphology, has been linked to a loss of channel inactivation (Bae et al., 2013), as well as increased channel sensitivity to mechanical force and increased trafficking (Glogowska et al., 2017). The net result in all these mutations is increased Ca^2+^ flux, and hence RBC dehydration. The loss of function phenotype is presumably due to a portion of variants not reaching the membrane (i.e. G2029R)(Lukacs et al., 2015)) and a proportion that are less sensitive to applied force.

Although lipids are of critical importance to all membrane embedded proteins (Laganowsky et al., 2014; Lee, 2004), they hold particular significance for mechanically-gated ion channels (Caires et al., 2017; Eastwood et al., 2015; Moe and Blount, 2005; Nomura et al., 2012; Perozo et al., 2002). This is because in many cases, such as the prototypical bacterial channels (Sukharev, 2002; Sukharev et al., 1994), mechanically-gated channels can sense forces directly from the membrane (Brohawn et al., 2014; Dong et al., 2015; Murthy et al., 2018). Akin to the bacterial mechanically-gated channels, Piezo1 has been shown to function in ‘reduced systems’ such as reconstituted bilayers (Cox et al., 2016; Syeda et al., 2016). This suggests the channel is not completely reliant on direct cytoskeletal connections. This in no way precludes a role for the cytoskeleton in Piezo1 function (Gottlieb et al., 2012; Poole et al., 2014), as membrane forces are largely determined by the local arrangement of the cytoskeleton and extracellular matrix. However, these experiments highlight that lipids are of critical importance to Piezo1.

Cryo-EM structures of the trimeric assembly of mouse Piezo1 from three separate groups show a triskelion arrangement and a peculiar cup-shaped topology (Guo and MacKinnon, 2017; Saotome et al., 2018; Zhao et al., 2018). The long arms (‘propellers’) of each monomer extend out and curve towards the extracellular space, with a long helix termed the ‘beam’ running almost parallel to the intracellular side of the propeller (Figure 1A). The propellers converge on a central pore region created by the last two transmembrane helices in the C-terminus, which sits under a conserved cap domain. The propellers are separated from the pore by an ‘anchor domain’, which forms a triangle and consists of two elbows and a helix that sits parallel at the membrane interface. In a simplified liposomal system, the channel itself seems to locally deform the membrane, retaining the cup-shape seen in all three structures (Guo and MacKinnon, 2017). While functional data points to a role for cholesterol (Qi et al., 2015), phosphatidylinositol 4,5-bisphosphate (PIP_2_) (Borbiro et al., 2015) and lysophosphatidylserine (Tsuchiya et al., 2018) in mechanical gating, exactly how lipids interact with Piezo1 channels and whether local curvature induced by the protein is retained in complex cellular systems is yet to be explored.

**Fig 1.**
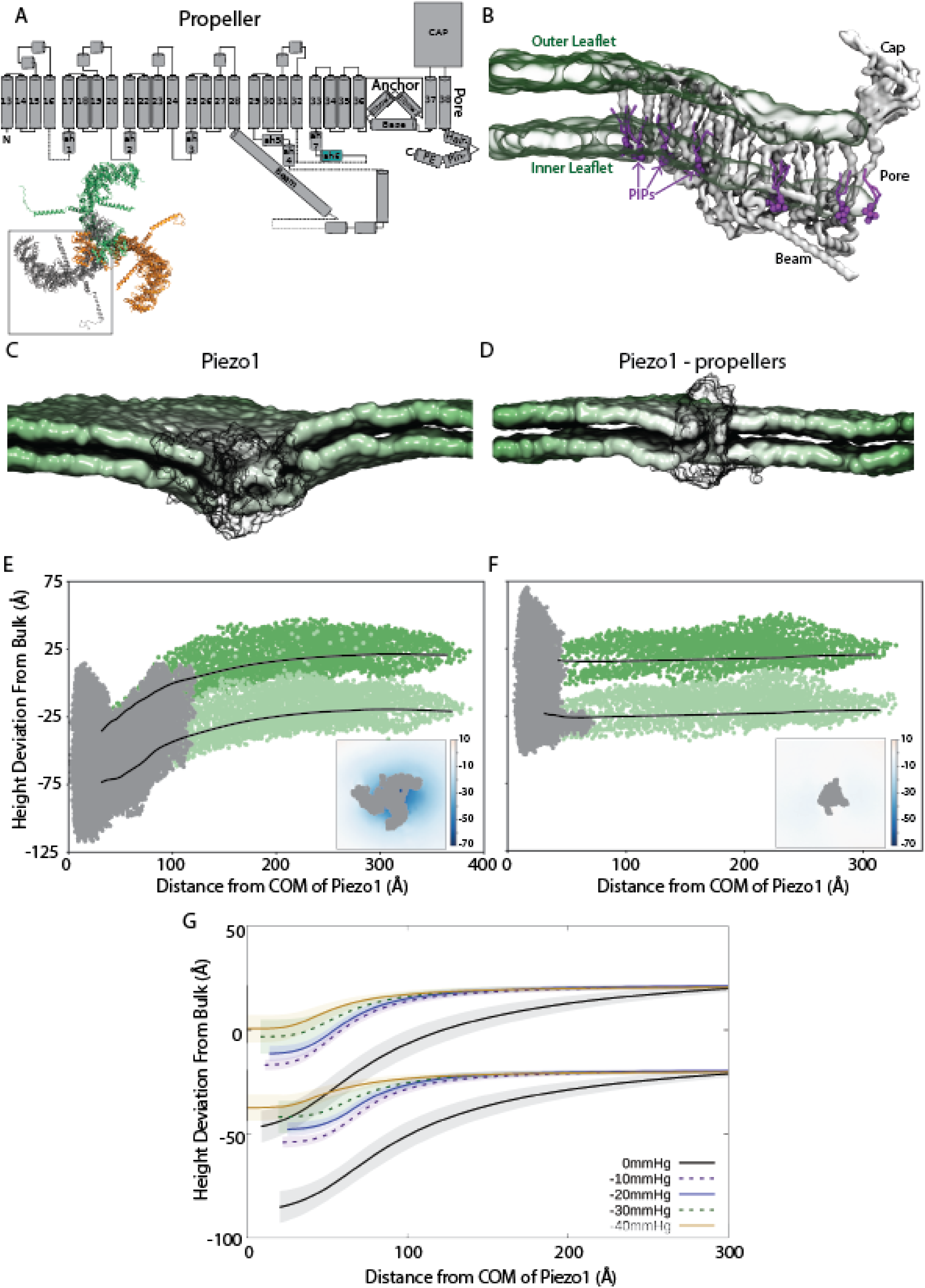
Local curvature induced by Piezo1 is driven by its propeller domains. (A) Schematic of the helix arrangement within each Piezo1 monomer, with the Piezo1 trimer shown below. (B) A side-on view of one of the propeller domains and the curved bilayer from simulations. Phosphate atoms are shown in green glass, Piezo1’s monomer is shown in grey surface, and phosphoinositides (PIP) lipids are shown in purple. Representative images of the mouse Piezo1 simulation systems with (C) and without (D) the propeller domains. The protein is represented in black glass, and the lipid phosphates are represented by the green surfaces. Lipid tails, water and ions are omitted for clarity. The average height of each membrane leaflet (as measured by the position of the phosphate atoms) as a function of distance from the centre of mass (COM) of the pore of Piezo1 with (E) and without (F) propellers (black lines in both graphs). Grey dots represent the average Cα atoms of Piezo1 and the dark and light green dots are the positions of the phosphate atoms of the top and lower leaflets of the last frame of each simulation system. The average height of the bilayer (in Å) as seen looking down onto the membrane during simulations of Piezo1 is shown in the insets. (G) Average height of the membrane in simulations under lateral tension. Each line represents the average of three repeat simulations, with the standard deviation shown as a shaded region above and below each line.

## Results

### Piezo1 induces significant local curvature in complex mammalian bilayer mediated via its propeller domains

To probe the local interactions of Piezo1 with surrounding lipids, we made use of coarse-grain molecular dynamics (CG-MD) simulations, an efficient tool to study the molecular details of protein induced membrane deformations (Ingolfsson et al., 2016). To closely mimic the composition of mammalian membranes, we simulated Piezo1 inside a complex asymmetric *in silico* bilayer containing more than 60 different lipid types (Ingolfsson et al., 2014). During each of three replicate 30 μs simulations of a ‘full-length’ construct of Piezo1, the membrane rapidly curves around the protein (Figure 1B,C). This protein model contains all regions of the protein resolved in the recent cryo-EM structures (pore, cap and beam domains plus six of the nine four-helical repeats that make up the propellers – see Figure 1A). The simulations begin with a flat bilayer, but the curvature develops rapidly, starting from the first frame of the simulation and stabilising within 1 μs. (Figure S1, top). This local curvature is maintained throughout the simulation, as indicated in Figure 1E, which shows the average height of each leaflet as a function of the distance from the centre of the protein over the last 10 μs of the 30 μs simulation. The large degree of membrane curvature is contained locally around Piezo1, and does not extend much beyond the ends of the propeller domains. This finding is corroborated by plotting the average height of the membrane as viewed from the extracellular space (inset of Figure 1E), which shows a very large degree of membrane curvature close to the protein. The curvature induced by Piezo1 is far greater than any previously simulated integral membrane protein (Corradi et al., 2018).

While the lipid bilayer undulates throughout the simulation, the curvature seen is not due to the chosen setup, nor the inherently asymmetric lipid composition of the bilayer: it is due to Piezo1’s propeller domains. This is demonstrated by three replicate simulations of a protein model lacking the propeller domains (Piezo1 – propellers), in which the membrane remains flat, even close to the protein (Figure 1D, F, S1). This reduced protein construct contains the pore domain and only one of the nine four-helical repeats in the propeller domains. Previous work suggests that a covalent link between the N and C terminal halves of Piezo1 is not necessary for mechanical gating of Piezo1 (Bae et al., 2016), but both halves have to be present. This is a finding we have replicated in Piezo1 KO cells, suggesting the importance of the propellers for mechanosensation. Interestingly, the thickness of the bilayer differs from that in bulk near the propeller domains (Figure S2). This is not seen in simulations without the propeller domains, reinforcing the idea that Piezo1 affects its local lipid membrane environment, predominantly via the propeller domains.

To compare the magnitude of curvature seen in our simulations with experimental observations, we first determine how the membrane curves along the surface of each single propeller domain as shown in Figure 1B. Here it can be seen that both the upper and lower leaflets follow the shape of each propeller domain, with the angle of the beam relative to the pore axis largely dictating the magnitude of curvature. In the cryo-EM structure obtained in a detergent micelle, Guo and Mackinnon calculate this angle to be ∼60° (where 90° represents flat propellers). In our simulations, this angle fluctuates between 50 and 65 degrees (Figure S3), in excellent agreement with the structural data. Guo and Mackinnon also proposed dimensions of the membrane curvature seen in their structure, based on the dome shaped deformation of the micelle surrounding Piezo1. The depth and radius of curvature of the dome is estimated as 60Å and 90Å, respectively. The depth is very similar to that seen in our simulations, where the difference between the average z coordinate of the bulk lipid and the average z coordinate of the lipid nearest the protein is almost exactly 60 Å (SI Figure 3B). The radius of curvature was determined using the second derivative of the average z coordinate, or concavity (Figure S3B), and yields approximately 122Å. While the radius of curvature in the simulations is larger than that seen in the micelle, it matches the curvature seen in a liposome (∼110Å), and can be overlaid closely upon the distorted liposome EM image of Guo and Mackinnon(Guo and MacKinnon, 2017). The dimensions of the curved footprint are expected to be larger in a bilayer than seen in the micelle or liposome (Haselwandter and MacKinnon, 2018). Interestingly, the curvature is not generated by the amphipathic helices in the propeller domains. Simulations in which these are deleted show greater flexibility in the protein and more membrane curvature (Figure S4). Notably, simulations under lateral membrane tension show a reduction in the membrane curvature in each of three repeat simulations. (Figure 1G). This is consistent with the hypothesis that force detection by Piezo1 can involve sensing changes to local membrane curvature. The curvature changes are greatest at low tension values, suggesting that sensitive detection of small forces may be enabled by this mechanism.

### Piezo1 induces curvature *in situ* in an intact cellular environment

The next question we aimed to address is whether the curvature seen in our simulations and *in vitro* in simplified liposomal membranes by Guo and Mackinnon is a realistic depiction of what occurs in cells. In order to do this, we used both fluorescence and an electron microscopy (EM) involving the plant derived ascorbate peroxidase, APEX2 (Figure 2). This technique has previously been used to interrogate the localization and morphology of other curved membrane structures, most notably caveolae (Ariotti et al., 2015). Specifically, it provides high fidelity localization within the depth of the sample (unlike ‘on-section labeling) in combination with nanometer resolution. Here, we used two strategies to localize Piezo1. We first used GFP and mCherry-fused Piezo1 that have been previously characterized (Figure 2A, left), including stable HEK293T lines expressing these proteins (Cox et al., 2016; Maneshi et al., 2018). We noticed two striking features: 1) tortuous structures in the ER and other intracellular membranes (Figure 2E) and 2) membrane “pits” highly enriched in Piezo1 (Figure 2F). In order to ensure these were Piezo1 dependent, and to increase fidelity further, we generated a Piezo1-1591-APEX2 fusion protein (Figure 2A, right) which functioned similar to WT, including retaining sensitivity to the small molecule agonist Yoda-1 (Figure 2B,C,D). Both the APEX2 fusions and GFP fusions show the same behaviour in EM micrographs when transiently expressed in HEK293T Piezo1 KO cells. The tortuous structures in the ER are particularly noticeable but are likely an artefact of over expression. Nevertheless, they provide evidence that Piezo1 in a cellular environment can generate profound curvature and bend biological membranes. The Piezo1 enriched membrane pits are between 90-150 nm in diameter, which is considerably larger than the curvature induced by an individual protein (d = ∼20-25 nm). The membrane pits are significantly larger than caveolae (Richter et al., 2008) but exhibit some morphological similarities (e.g. like caveolae they can also be observed in chains (Figure 2G)). In order to examine their possible relationship to caveolae we expressed Piezo1 in caveolin-1 null mouse embryonic fibroblasts that completely lack caveolae (Kirkham et al., 2008). Piezo1 assocaited with morpholically identical surface pits in cells lacking caveolae. In addition, Piezo1 associated with complex internal curved structures (Figure 2H-I) further arguing that these curved structures are Piezo1-dependent.

**Fig 2.**
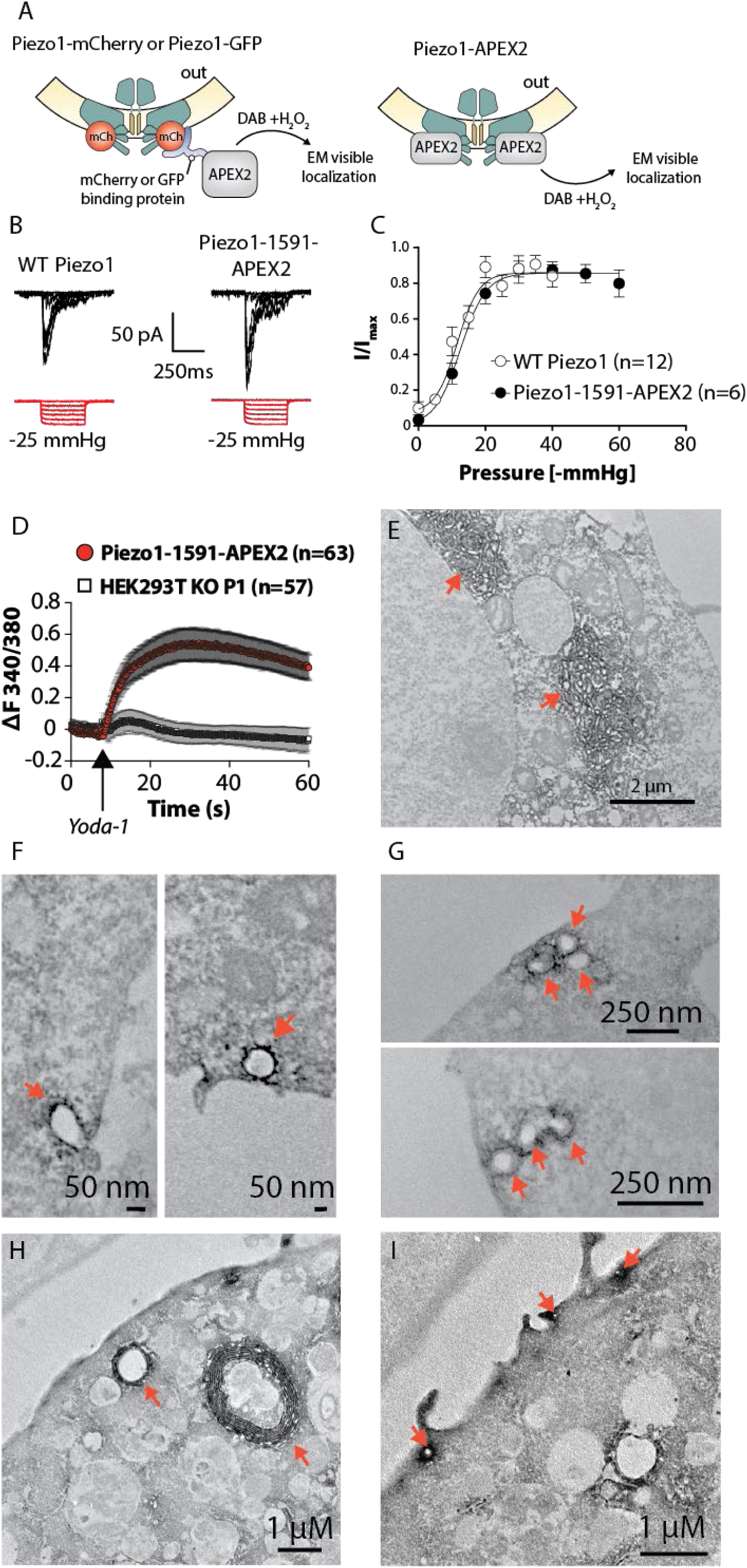
Electron microscopy utilizing APEX2 reveals cellular curvature induced by Piezo1 expression. (A) Pictorial description of the strategy used to determine whether Piezo1 induces local curvature. We used the plant ascorbate peroxidase (APEX2) fused to a GFP or mCherry binding protein (left panel) or a direct Piezo1-1591-APEX2 fusion protein. (B) Representative traces of Piezo1 activity and Piezo1-1591-APEX2 evoked by negative pressure pulses at a holding potential of −65mV. (C) Comparison of pressure sensitivity between WT Piezo1 and the Piezo1-1591-APEX2 fusion protein. (D) Ratiometric Fura-2 Ca^2+^ imaging in Piezo1-1591-APEX2 expressing cells and Piezo1 KO HEK293T. Arrow indicates the timepoint at which 1 µM Yoda-1 was added. (E) EM image showing tortuous ER structures from Piezo1-1591-APEX expressing cells (using Piezo1 KO HEK293T as the background). These structures are never seen in non-transfected controls. (F) Examples of curved pit-like structures at the plasma membrane of between 90-150 nm in diameter densely stained with Piezo1. (G) Chains of ‘Piezo1 pits’ at the plasma membrane somewhat reminiscent of the chains of structures created by caveolae. (H) Intracellular curved structures in Cav-1 KO mouse embryonic fibroblasts expressing Piezo1-1591-APEX2. (I) Dense staining of Piezo1-1591-APEX2 in membrane invaginations in Cav-1 KO mouse embryonic fibroblasts.

Given their size, and the density of Piezo1 staining, these structures must involve multiple Piezo1 timers and can plausibly be explained by the cooperative curvature induced by multiple neighbouring Piezo1 proteins akin to that induced by other triskelions such as clathrin (Haucke and Kozlov, 2018) and oligomeric integral membrane proteins such as caveolin (Stoeber et al., 2016) and synaptophysin (Adams et al., 2015). This is supported by the fact that Piezo1 proteins are known to accumulate together as clusters (Bae et al., 2013; Ridone et al., 2019).

### Piezo1 forms functionally important, localised interactions with specific lipids

In addition to the local curvature and thickening of the bilayer induced by Piezo1, the CG-MD simulations reveal enrichment and depletion of specific lipids around the protein, suggesting a clear preference for the protein to interact with some lipids over others (denoted lipid fingerprints (Corradi et al., 2018)). Some lipid types are highly enriched around the protein, in particular glycolipids, phosphatidylinositolphosphates (PIPs), and diacylglycerols (DAGs); while some lipids, most notably sphingomyelin (SM), are strongly depleted. This can be inferred from the Depletion/Enrichment (D/E) index calculated for each lipid group for each leaflet (Figure 3A), as well as by density plots of specific lipids around the protein (Figure 3B) and the residue-lipid contact maps (Figure S5 and S6). The lipid fingerprint of Piezo1 is similar to that observed for other membrane proteins (Corradi et al., 2018), including the relative higher occupancy of polyunsaturated lipids compared to saturated ones. As seen in Figures S5 and S6, a large number of lipids including phosphatidylserines (PS), phosphatidic acids (PA), SM, phosphatidylethanolamines (PE), PIPs, phosphatidylinositols (PI) and cholesterol bind at specific sites on the protein. This includes some lipids that are not enriched overall in the neighbourhood of Piezo1.

**Fig 3.**
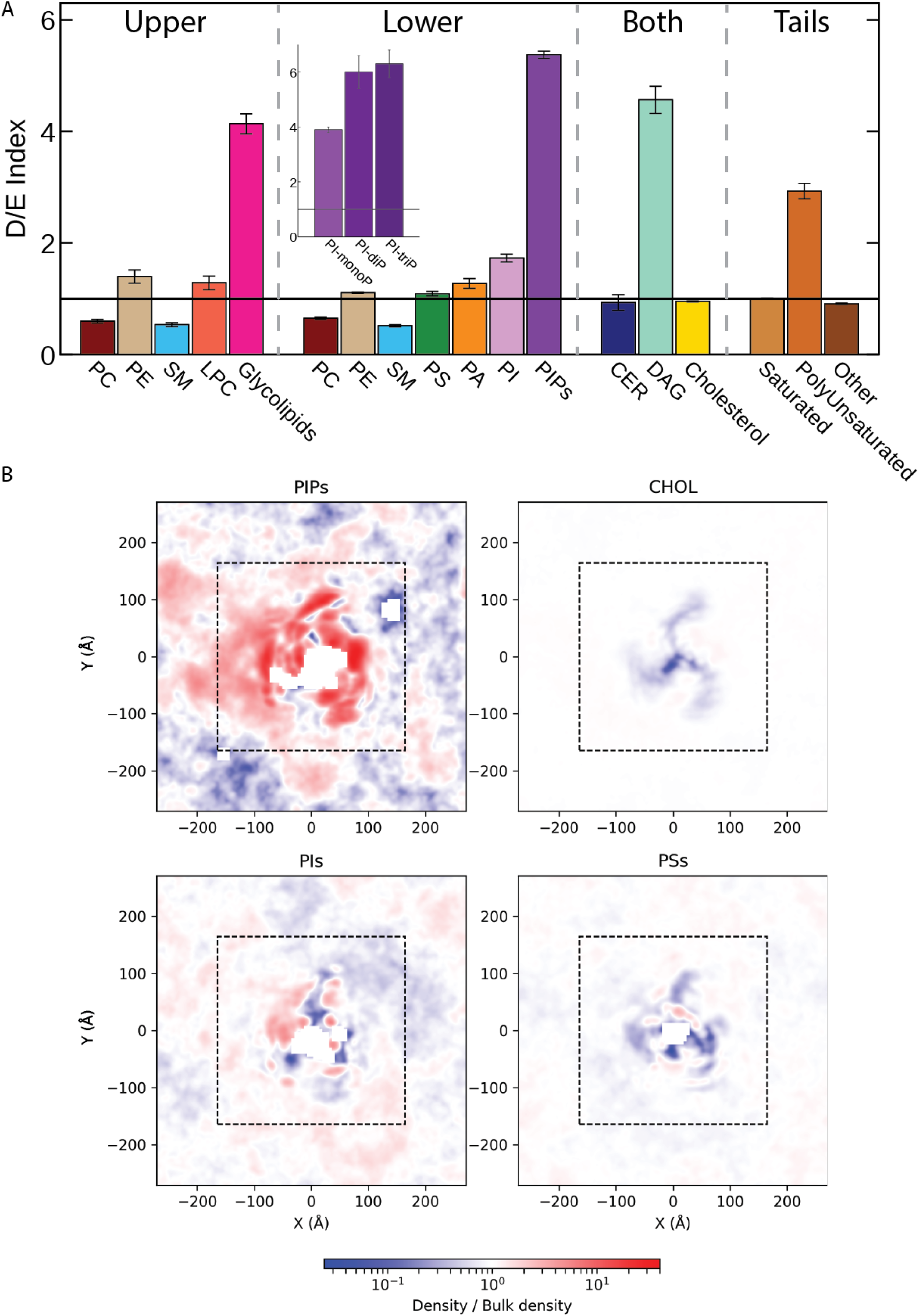
Lipid fingerprinting of the Piezo1 channel. (A) Average and S.E.M. of the Depletion/Enrichment (D/E) index of all lipid groups in three simulations within a 20Å cut off of Piezo1. Lipids are separated by leaflet, and by their tendency to flip-flop (these are noted as ‘both’). The D/E index is also calculated for lipids grouped according to tail saturation. D/E of different PIP species are shown in the inset. (B) Densities of PIPs, Cholesterol, PIs, and PS/PAs around Piezo1, shown as a fraction of the bulk density.

A lipid class of particular interest is PIPs, which includes not only the aforementioned PIP_2_, but also a range of other PI-monophosphates, PI-diphosophates and PI-triphosphates (noting that CG-MD does not distinguish within these categories). PIPs (especially the di- and tri-phosphate species) are not only highly enriched around the protein (inset to Figure 3A) but can be seen to form high-density regions around both the pore domain and propellers (Figure 3B). The clustering of PIPs to specific sites is also evident by looking at selected snapshots, as depicted in Figure 1B and the inset of Figure 4A. To quantify this further, we calculate which residues of the protein most frequently interact with PIPs, and map this on to the protein structure in Figure 4A. Among many specific protein-lipid interactions identified, an intriguing binding site for PIPs was determined around the pore domain, shown on the right in Figure 4A. There is a noticeable patch of four lysines, K2166-K2169 (human sequence numbering), right before helix 37, which is highly conserved in Piezo1 channel homologues (Figure 4B). These lysines, along with some adjacent lysines (K2096-K2097, and K2163), were identified to consistently interact with PIPs (most notably di-phosphate species, Fig S7), as seen by the constancy of protein-PIP contacts at these sites (Figure 4C). To test the importance of this site, we removed this string of four lysines (Δ4K mutant), as this variant had been noted to cause xerocytosis (Albuisson et al., 2013). Patch clamp electrophysiology in the cell-attached configuration show that this mutation reduces channel inactivation significantly. Using large cell-attached patches with ‘mesoscopic’ currents arising from multiple channels, it was not possible to measure the inactivation rate because the non-inactivating phenotype was so severe (Figure 4D,F), although the pressure sensitivity of the mutant was unchanged from WT (Figure 4E). As previously described (Bae et al., 2013), we measured the degree of steady state current, which clearly quantifies the marked difference with WT (Figure 4G). We also used depolarising voltages to compare deactivation, and again deactivation of the Δ4K mutant was slower than WT (Figure 4H-J). Simulations of this mutant show a large reduction in PIP enrichment around the protein, and specifically a loss of mono- and di-phosphate PIP molecules at this site (Figure S7, top row). A mutation of the residue K2097E (K2113E in mouse) also has a modest effect in speeding up inactivation, though probably not by preventing PIP binding (Figure S7, middle row).

**Fig 4.**
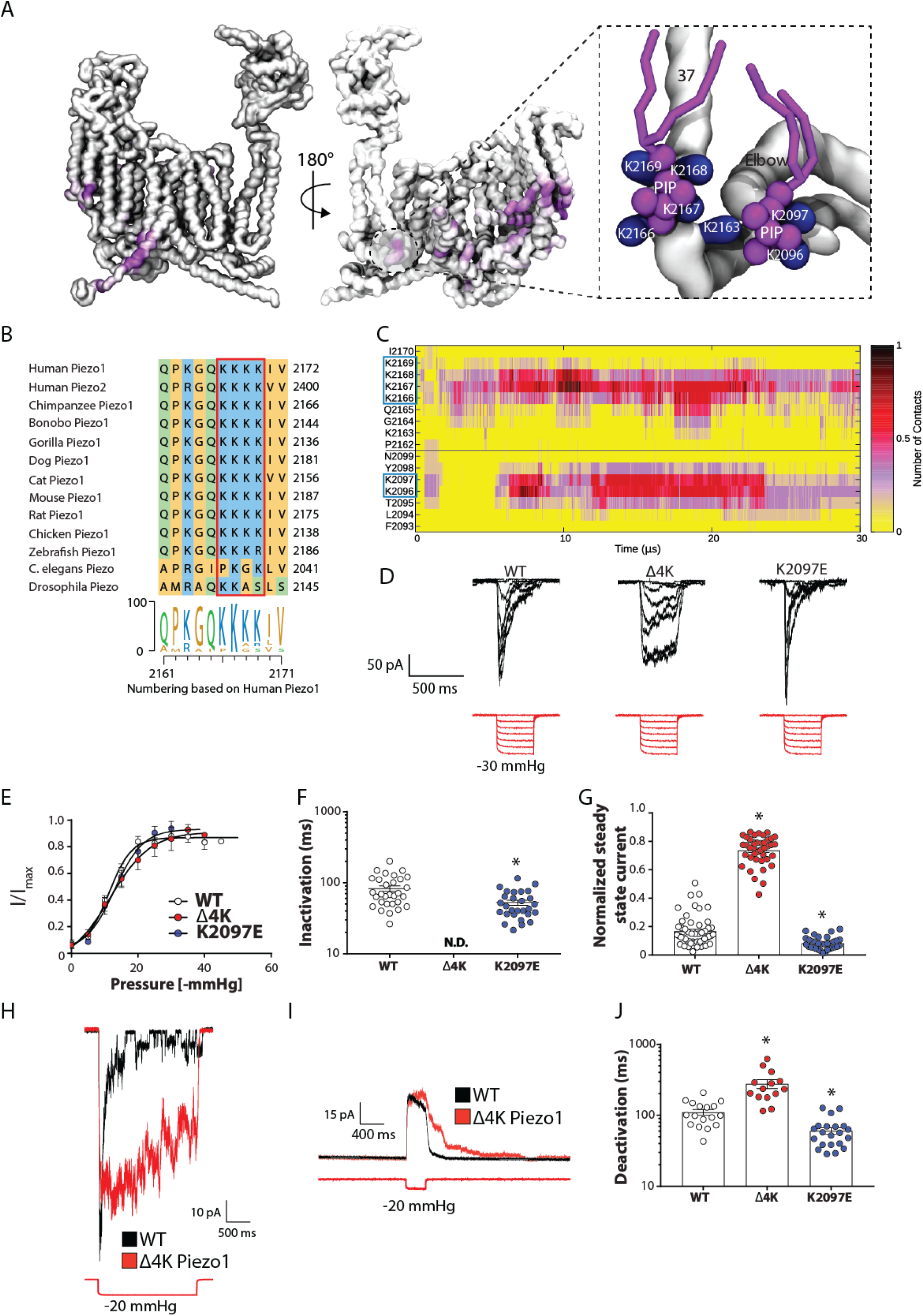
Computationally identified Phosphatidylinositide binding sites and the effect of mutations on Piezo1 channel function. (A) Contact map of a Piezo1 monomer indicates residues that form consistent interactions with PIPs (purple) with the protein backbone represented as a silver surface. Inset shows a snapshot of a PIP binding site close to the pore, with relevant lysines shown in blue and PIPs in purple. Residue numbering corresponds to the human sequence for comparison with experimental data, although simulations are conducted on mouse Piezo1. (B) Complete conservation of human Piezo1 K2166-2169 in mammalian homologues. (C) Contact analysis showing interactions between PIPs and Piezo1 residues, highlighting the stability of the interaction of specific residues with PIPs throughout the simulation. (D) Deletion of conserved lysines in human Piezo1 is associated with xerocytosis and causes a loss of inactivation. Representative traces of WT human Piezo1, mutant where these conserved lysines are deleted (Δ4K) and charge reversal at another site located near these conserved lysines (hK2097E, equivalent to mK2113). All cell-attached traces recorded at a holding potential of −65mV. (E) Pressure response curves of WT Piezo1 and the two mutants (n=>6). (F) Inactivation time constant in ms comparing WT to the PIP binding mutants from pressure pulses of 20-40 mmHg. The inactivation phenotype of Δ4K mutant was so profound that accurate fitting of ‘mesoscopic’ cell-attached currents was not possible. (G) Analysis of steady state current from pressure pulses of 20-40 mmHg show a large increase in Δ4K and a modest but statistically significant decrease in the K2097E mutant. Data represents mean ± S.E.M. (H) Long 2s square wave pressure pulse illustrating the vast difference in the Δ4K mutant to WT Piezo1. (I) Representative current traces from cell-attached patches of PIP binding mutants in response to a square wave pressure pulse at +90 mV holding potential. (J) Quantification of Piezo1 deactivation at +90 mV holding potential.

Our simulations also show that R808 binds all species of PIPs (Figure S7, bottom row). R808Q is a rare variant with a minor allele frequency of 0.6% and has been reported as a gain of function mutant associated with xerocytosis (Andolfo et al., 2013). However, this mutation has been reported in patients in combination with G782S, though which of these are disease causing has yet to be addressed. Simulations of the R808Q mutant show a loss of PIP binding but no loss of other lipids (Figure S7, bottom row), suggesting that the gain of function may be related to changes in the pattern of lipid binding.

Previous work suggested R2119 is essential and may interact with negatively charged lipids (Saotome et al., 2018). However, we did not see any negatively charged lipids interacting with this residue. We do, however, see a salt bridge between R2119 and E2140 that is present most of the time in all three monomers, and in all three replicates (Figure S8). This salt bridge may be the reason for the need for arginine at this position.

Since cholesterol is present as 30% of the membrane fraction in our CG-MD simulations, there are naturally a large number of residues on Piezo1 that interact with cholesterol (Figure S5). However, there are certain places on the structure of Piezo1 where cholesterol is found most frequently (Figure 5A) that may help understand the importance of cholesterol to Piezo1 function. We searched for cholesterol recognition motifs (termed CRAC and CARC)(Di Scala et al., 2017; Posada et al., 2014) within the sequence of Piezo1 using ProSite (https://prosite.expasy.org/scanprosite/), and identified 19 CRAC motifs, and 39 CARC motifs (Table S2). Previous crosslinking studies showed an interaction of cholesterol with four residues (LVPF), which are part of a CARC motif we have identified, and are located in the anchor domain between the two elbow regions of Piezo1 (Hulce et al., 2013), a region previously identified as being highly conserved (Prole and Taylor, 2013). Cholesterol was observed in this same site in all 3 monomers on average 75 ± 20% of the simulation duration (Figure 5B, bottom right), highlighting the ability of the coarse grained simulations to reproduce known protein-lipid interactions. To test the importance of predicted cholesterol binding sites, we examined a number of alanine mutations in this conserved region (i.e. P2113A) and in other parts of the first Piezo1 repeat as identified in simulations and by recognition sequences to interact with cholesterol (L1966A, L2037A & Y2093A) (Table S2, highlighted in yellow). L1966A and L2037A behave similar to WT with a similar P_1/2_ and inactivation properties (Figure 5C-E). However, P2113A has a right shifted pressure response curve (WT 11.92 ± 0.81, n=15; P2113A 20.18 ± 1.92 mmHg, n=15) (Figure 5D). Y2073A could not be opened regardless of the pressure applied up to the lytic limit of the patch. Recent work suggests that another residue that forms a cholesterol recognition site (Y2022H) is also a loss of function mutation resulting in bicuspid aortic valve (Faucherre et al., 2019).

**Fig 5.**
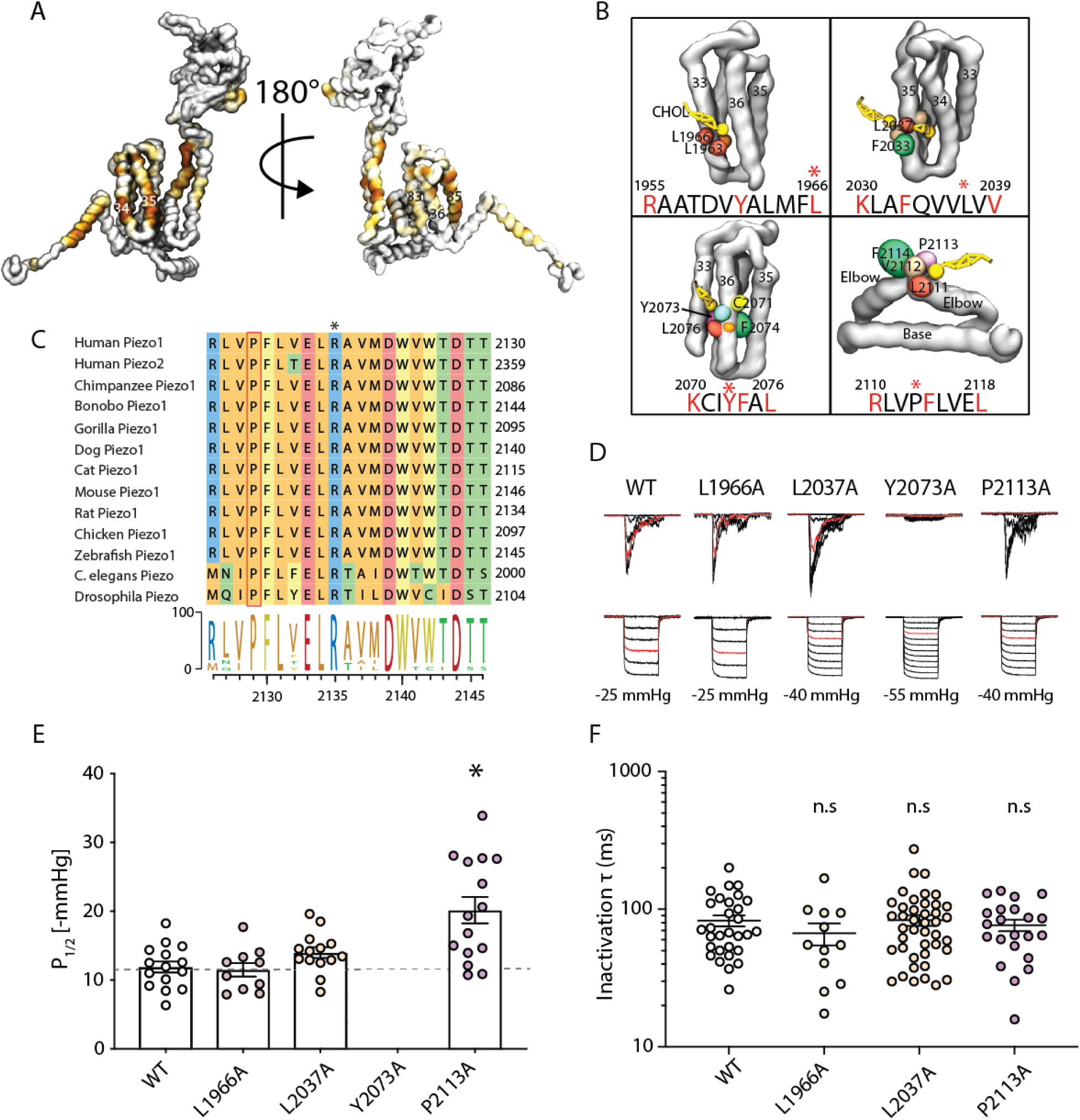
Computationally identified cholesterol binding sites and the effect of point mutations on Piezo1 activation. (A) Cholesterol (yellow) contact map of a Piezo1 monomer (only the regions close to the pore shown for clarity) with the backbone represented as a silver surface. Brown regions denote areas where a cholesterol molecule resides greater than 65% of the simulation. (B) Snapshots of four CARC motifs near the pore domain that bind cholesterol. The relevant sequence and relevant residues are highlighted, and the human numbering is shown. Residues that were subsequently mutated are indicated by *. (C) Alignment of anchor domain and conservation of human Piezo1 P2113 in all Piezo1 homologues. (D) Cell-attached currents of human Piezo1 WT and residues identified to interact with cholesterol; L1966A, L2037A, Y2073A and P2113A. All are part of bona fide cholesterol recognition motifs. Currents recorded at a holding potential of −65 mV and for comparison the −15 mmHg pulse of each representative trace is shown in red. (E) The calculated P_1/2_ from each patch compared to WT. (F) Inactivation constant from cell-attached patches showing that there is little difference between these mutants and WT. Data are means +/- S.E.M * represents p<0.01.

### Bidirectional feedback between Piezo1 and membrane lipids

We have already shown that Piezo1 locally distorts the membrane and interacts with specific lipid types. Given that lipids affect Piezo1 activity, it is possible that there is bidirectional feedback between the expression of Piezo1 and lipids. To probe this question, we used array-based transcriptomics to identify lipid-regulating proteins that were differentially expressed in the presence or absence of Piezo1. We compared the expression of lipid-regulating proteins in Piezo1 KO HEK293T cells with two over expressing lines, each with a graded increase in Piezo1 expression (Fig 6A). There is a large change in a number of genes associated with lipid regulation (Fig 6B-C). The first striking difference we noted was in N-acylethanolamine (NAE) metabolism. Physiologically important NAEs, such as palmitoylethanolamide and anandamide, are essential in vascular biology (Montecucco and Di Marzo, 2012), as is Piezo1 (Li et al., 2014; Ranade et al., 2014). There is a vast Piezo1-dependent difference in N-acylethanolamine acid amine hydrolase (NAAA1) and fatty acid amide hydrolase 2 (FAAH2) expression levels (FAAH2 6.8 and 8.2-fold down compared to KO, NAAA 16.0 and 22.7-fold up compared to KO (representing the extreme ends of Fig 6C). We confirmed this differential expression using qPCR (Fig 6C,D). FAAH2 and NAAA are ubiquitous enzymes that break down N-acylethanolamines of different types, and have a differential localization and preference for NAEs (Tsuboi et al., 2005). The fact that the NAE-digesting enzymes FAAH2 and NAAA have large changes in expression, and the precursors of NAEs (which are N-acyl phosphatidylethanolamines (NAPEs)) are formed by Ca^2+^-dependent *N*-acyltransferases, means Piezo1’s transport of Ca^2+^ ions through mechanical stimuli can input a relevant stimulus into the NAE signalling pathway.

**Fig 6.**
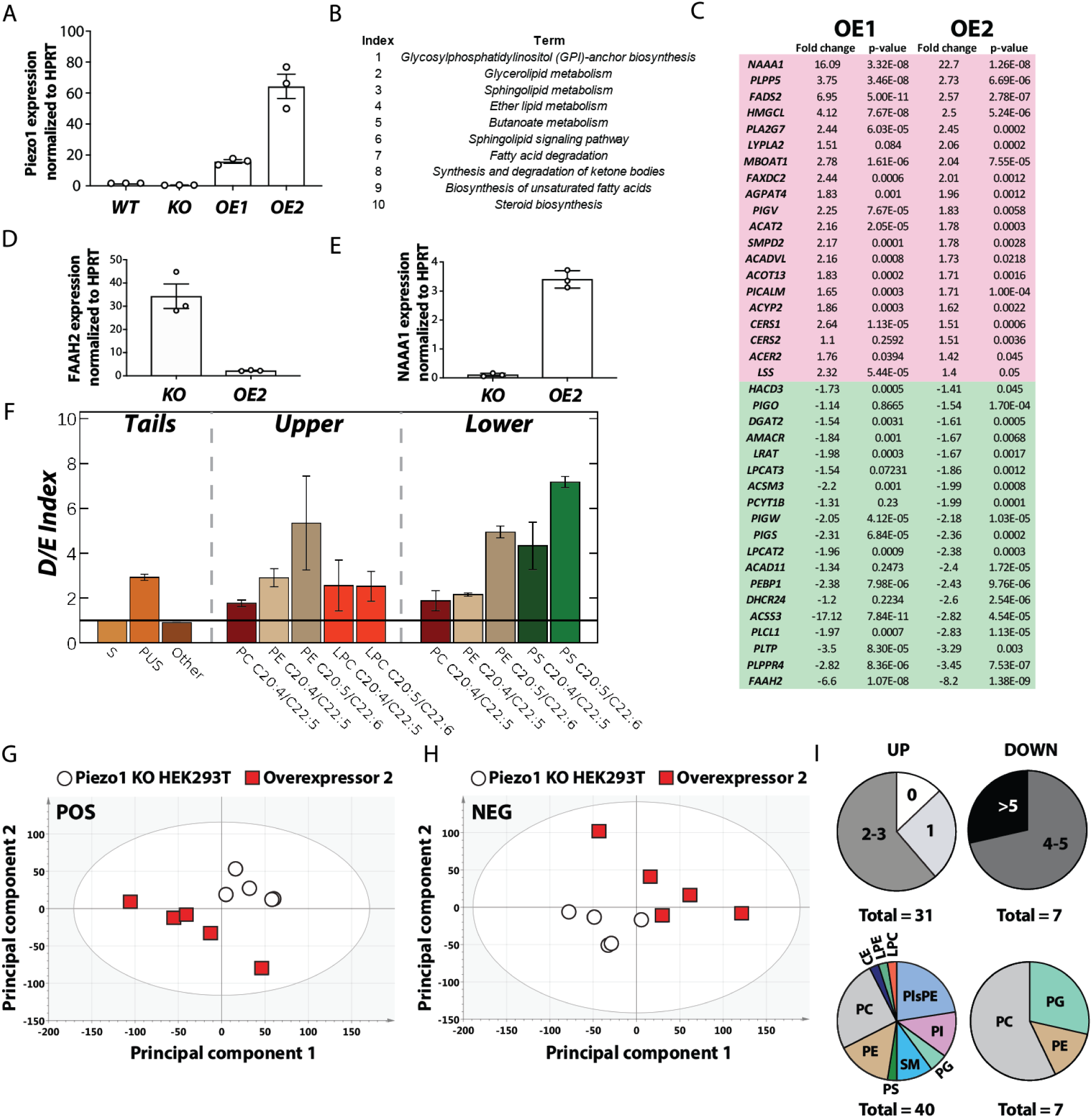
Piezo1 expression grossly affects lipid metabolizing networks and the cellular lipidome. (A) Relative expression of Piezo1 in cell lines used in this study [KO – Piezo1 KO HEK293T, WT – HEK293T, overexpressor 1 “OE1” – Piezo1-1591-GFP HEK293T, overexpressor 2 “OE2” - Piezo1-1591-mCherry]. (B) Table illustrating genes pathways affected by overexpression of Piezo1. (C) Table illustrating genes involved in lipid regulation modified in Piezo1 overexpressing cells compared with the KO. (D) Validation of differential expression of FAAH2 using qPCR. (E) Validation of differential expression of NAAA1 using qPCR. (F) *In silico* effect on local lipidome in the form of an average depletion/enrichment (D/E) index of specific polyunsaturated lipids, shown with S.E.M. (G) PCA scores plot of positive ionisation data from multivariate analysis of the lipidome from KO vs OE2 [Piezo1-1591-mCherry] (2 components; R^2^[cum] = 0.490; Q^2^[cum] = 0.170). (H) PCA scores plot of negative ionisation data from KO vs OE2 [Piezo1-1591-mCherry] (2 components; R^2^[cum] = 0.479; Q^2^[cum] = 0.152). (I) Graphical representation of the changes seen in lipids in the OE2 [Piezo1-1591-mCherry] cell line compared with KO HEK293T. Upper panel shows changes in saturation quantified as the number of double bonds per lipid class that is differentially expressed and lower panel shows the lipid classes that are modified by Piezo1 expression.

Our simulations and previous work (Romero et al., 2019) point to a critical role in acyl chain saturation in modulating Piezo1 function. Transcriptomics show fatty acyl-coenzyme A (CoA) desaturase (FADS2) is 6 and 3-fold up-regulated in Piezo1 over-expressing cells compared to KO. This enzyme acts to introduce double bonds at carbon 6 in fatty acid chains, and is particularly important in the synthesis of polyunsaturated fatty acids. Other lipid regulating enzymes differentially regulated in response to Piezo1 are shown in Fig 6C.

Finally, we sought to address whether these changes in lipid regulating enzymes culminated in Piezo1-dependent changes in the whole cell lipidome. The HEK293T KO and OE2 (overexpressor 2) cell lines were found to have significant differences in their lipidomic profiles (by Student’s t-test) in both positive and negative ionisation mode. Three different analysis methods showed many thousands of lipid components to have changed between the two cell lines (see STAR methods), however only those that could be identified with the highest level of assurance are discussed below.

Multivariate analysis of XCMS-processed data using principal component analysis shows a clear separation between the two cell lines for both ionisation mode datasets (Fig 6G-H), which are further resolved by using a supervised method (orthogonal partial least squares-discriminant analysis; OPLS-DA) to partition the variance into predictive and orthogonal (unrelated) components (Fig S9, A-B). Both OPLS-DA models were validated via permutation (x 100), with permutation plots showing the requisite predictive failure of randomly permuted models (Fig S9 C-D). The cell line-dependent changes observed in the lipidome summarised in Fig 6I and detailed in Table 1 show alterations in several lipid classes (CE, [L]PC, [L]PE, PI, PG, SM, as well as plasmenyl/plasmanyl PE species). Although class-specific changes were observed in both directions, there was a trend towards more saturated species in the over-expressor compared to the knockout cell line. Recent data show that saturated fatty acids increase membrane bending stiffness and inhibit Piezo1, while polyunsaturated fatty acids can decrease membrane stiffness and modulate inactivation (Romero et al., 2019). Our simulations show that polyunsaturated lipids accumulate near Piezo1 (Fig 3A, 6F), presumably to aid in generating membrane curvature. The trend toward more saturation in cells overexpressing Piezo1 may represent an attempt to reduce Piezo1 activity, although the picture is clearly nuanced with a variety of changes in lipid saturation. The over-expressing cell line also exhibited increased abundance in several plasmenyl/plasmanyl PE lipids, as well as a cholesteryl ester. Plasmenyl lipids contain a vinyl ether and have previously been shown to modify and stabilise cellular structures of profound negative curvature such as caveolae and clathrin coated pits (Thai et al., 2001). Furthermore, an increase in cholesteryl ester fits with the corresponding rise measured in ACAT2, an enzyme which catalyses the conversion of cholesterol to its corresponding esters.

**Table 1.** List of identified changes in the lipidome of Piezo1 overexpressing cells. [KO vs OE2 Piezo1-1591-mCherry]. Results are ordered from most upregulated (pink) to most downregulated (green).

| Species | Class | Fold Change | p-value | Ionisation |
| --- | --- | --- | --- | --- |
| PC 26:0 | Phosphatidylcholine | 5.53073 | 0.0115 | POS |
| PC 28:1 | Phosphatidylcholine | 5.27023 | 0.0110 | POS |
| Plasmanyl-PE(O-36:4) | Plasmanyl-phosphatidylethanolamine | 4.8778 | 0.0222 | NEG |
| PC 27:0 | Phosphatidylcholine | 4.84285 | 0.0153 | POS |
| PC 30:2 | Phosphatidylcholine | 4.70117 | 0.0178 | POS |
| PC 20:1 | Phosphatidylcholine | 4.3249 | 0.0107 | POS |
| PE 38:3e | Phosphatidylethanolamine | 4.19552 | 0.0447 | POS |
| CE 18:1 | Cholesteryl Ester | 3.78796 | 0.0000 | POS |
| PI 40:2 | Phosphatidylinositol | 3.78759 | 0.0117 | NEG |
| Plasmenyl-PE(P-37:2)) | Plasmenyl-phosphatidylethanolamine | 3.70895 | 0.0174 | NEG |
| LPC 14:0 | Lysophosphatidylcholine | 3.66381 | 0.0057 | NEG |
| PE 32:2e | Phosphatidylethanolamine | 3.63726 | 0.0083 | POS |
| Plasmenyl-PE(P-32:2)) | Plasmenyl-phosphatidylethanolamine | 3.63427 | 0.0173 | NEG |
| SM d40:2 | Sphingomyelin | 3.42926 | 0.0059 | POS |
| Plasmanyl-PE(O-38:3) | Plasmanyl-phosphatidylethanolamine | 3.364 | 0.0099 | NEG |
| PE 38:2 | Phosphatidylethanolamine | 3.28495 | 0.0061 | NEG |
| PE 40:2 | Phosphatidylethanolamine | 3.0288 | 0.0082 | NEG |
| LPE 18:1 | Lysophosphatidylethanolamine | 3.02437 | 0.0247 | NEG |
| PS 32:0 | Phosphatidylserine | 2.97411 | 0.0080 | NEG |
| PG 38:2 | Phosphatidylglycerol | 2.93944 | 0.0113 | NEG |
| Plasmenyl-LPE(P-18:1) | Plasmenyl-lysophosphatidylethanolamine | 2.92386 | 0.0216 | NEG |
| PG 38:3 | Phosphatidylglycerol | 2.91235 | 0.0047 | NEG |
| PE 30:1e | Phosphatidylethanolamine | 2.75658 | 0.0003 | POS |
| Plasmanyl-PE(O-40:3) | Plasmanyl-phosphatidylethanolamine | 2.75289 | 0.0265 | NEG |
| Plasmanyl-PE(O-36:3) | Plasmanyl-phosphatidylethanolamine | 2.75039 | 0.0098 | NEG |
| Plasmanyl-PE(O-34:3) | Plasmanyl-phosphatidylethanolamine | 2.58012 | 0.0152 | NEG |
| PI 40:3 | Phosphatidylinositol | 2.52118 | 0.0038 | NEG |
| PI 39:2 | Phosphatidylinositol | 2.51489 | 0.0146 | NEG |
| Plasmanyl-PE(O-31:1) | Plasmanyl-phosphatidylethanolamine | 2.38432 | 0.0013 | NEG |
| PI 37:3 | Phosphatidylinositol | 2.33399 | 0.0013 | NEG |
| PC 38:3 | Phosphatidylcholine | 2.30412 | 0.0012 | POS |
| PE 34:2 | Phosphatidylethanolamine | 2.13579 | 0.0077 | NEG |
| PI 36:3 | Phosphatidylinositol | 2.0332 | 0.0052 | NEG |
| SM 36:1 | Sphingomyelin | 2.02484 | 0.0066 | POS |
| PC 36:3 | Phosphatidylcholine | 1.91973 | 0.0065 | POS |
| PC 38:2 | Phosphatidylcholine | 1.69214 | 0.0014 | NEG |
| PC 42:5 | Phosphatidylcholine | -2.22051 | 0.0001 | POS |
| PC 40:5 | Phosphatidylcholine | -2.23044 | 0.0016 | POS |
| PE 36:5 | Phosphatidylethanolamine | -2.26285 | 0.0037 | NEG |
| PC 40:5e | Phosphatidylcholine | -2.42076 | 6.27E-05 | NEG |
| PC 34:5 | Phosphatidylcholine | -3.53378 | 0.000744 | POS |
| PG 44:12 | Phosphatidylglycerol | -3.92093 | 0.000593 | NEG |
| PG 44:11 | Phosphatidylglycerol | -4.01555 | 7.42E-07 | NEG |

## Discussion

### Piezo1 induces significant local curvature in complex mammalian bilayers mediated via its propeller domains

Here we have used a multi-disciplinary approach to interrogate the interaction of Piezo1 with its annular lipids and its propensity to induce local curvature *in silico* and *in situ*. Piezo1 supports local membrane curvature *in silico* in a complex mammalian membrane consisting of >60 lipid types. This curvature (*d* = ∼20 nm) is dependent on its propellers but not on the amphipathic helices associated with each Piezo1 repeat. Membrane curvature around the protein markedly decreases under lateral tension, consistent with the idea that Piezo can detect forces via alterations in local membrane curvature.

### Piezo1 induces curvature *in situ* in an intact cellular environment

Electron microscopy using a novel Piezo1-APEX2 fusion protein shows that the local curvature induced by an individual Piezo1 trimer (*d* = ∼20 nm) is magnified as the proteins cluster resulting in striking, larger curved structures (*d* = ∼120-150 nm) in cell membranes that we term ‘Piezo1 pits’. These are noticeable distinct membrane invaginations. While the membrane inherently undulates and contains both caveolae and other invaginations these ‘pits’ enriched with Piezo1 seem distinct in size and morphology and are present in cells devoid of caveolin-1. The exact role of these Piezo1 pits and the other structural elements which are likely involved should be the focus of future studies, but may represent a means of increasing the mechanosensitivity of the channels through enhanced local curvature (Haselwandter and MacKinnon, 2018) and cooperative behaviour.

### Piezo1 forms functionally important lipid interactions

Extensive analysis of lipid interactions identifies a characteristic lipid fingerprint for Piezo1. This includes enrichment of PIPs and phospholipids with unsaturated acyl chains. We identify a number of essential binding sites for phosphoinositides as well as cholesterol. Site-directed mutagenesis in combination with electrophysiology shows that these sites are functionally relevant and in many cases are related to Piezo1-mediated pathologies.

In particular, we identify a highly conserved PIP binding site consisting of a patch of 4 lysines which, when deleted, causes a xerocytosis phenotype (Albuisson et al., 2013). Analysis of this mutant channel shows a profound loss of inactivation. In addition, several other residues bind PIPs and are associated with disease, including R808Q.

Cholesterol has been shown to cross link with a number of regions of Piezo1 including the apex of the anchor domain, an area where cholesterol is enriched during our simulations (Hulce et al., 2013). Both this region (P2113A) and a cholesterol recognition motif (Y2073A) in the first Piezo1 repeat influence function. However many of the mutants are wild-type like, suggesting that cholesterol effects may be mediated by both specific protein lipid interactions and a global influence on the mechanical properties of the membrane (Cordero-Morales and Vasquez, 2018; Qi et al., 2015). Moreover, it is indeed interesting that lipid binding sites can have a positive or negative modulatory effect on Piezo1 channel function. These sites seem to influence not only mechanosensitivity but also gating kinetics. These results, in addition to recent work on specific and global effects of lipids on Piezo1, make it clear that Piezo1 mutations, where possible, should be studied in the membranes of natively expressing Piezo1 cells. Finally, the *in silico* and *in situ* curvature of the membrane induced by Piezo1 likely requires strong protein-lipid interactions to anchor the bilayer around the curved shape of Piezo1. More work is required to determine if the binding sites seen for specific lipid types are essential for curving the shape of the membrane.

### Piezo1 changes in the cellular lipidome

We also go one step further and show that Piezo1 causes changes in a host cell’s lipid synthesizing and metabolizing enzymes (e.g. NAAA1, FAAH2, ACAT2, FADS2 etc) and a corresponding change in the cellular lipidome. Given the profound changes in membrane shape that we see on overexpression using EM, and the recruitment of specific lipids seen in the simulations, it is unsurprising that the cell responds with modification of membrane lipid constituents. Such changes in the lipidome may act to tune Piezo1 activity or to stabilise membrane structures. Future work should aim at recapitulating these changes at physiologically relevant Piezo1 levels and deciphering whether Piezo1 mutations may differentially affect the local lipidome and in turn influence physiology and pathology.

## Acknowledgements

We thank Tsjerk Wassenaar for his invaluable advice and help with his Martinate script. We also thank Helgi Ingolfsson for usage of his original depletion/enrichment script, and for permission to publish it. We are grateful for funding from the Australian Research Council (FT130100781) that supported this research. SJM is supported by an ERC Advanced grant “COMP-MICR-CROW-MEM”. We grateful acknowledge the Victor Chang Cardiac Research Institute Innovation Centre, funded by the NSW Government, as well as funding from the Freedman Foundation for the Metabolomics Facility. This work was also supported by the National Health and Medical Research Council of Australia (grants APP1140064 and APP1150083 and fellowship APP1156489 to R.G.P.). RGP is supported by the Australian Research Council (ARC) Centre of Excellence in Convergent Bio-Nano Science and Technology. The authors acknowledge the use of the Microscopy Australia Research Facility at the Centre for Microscopy and Microanalysis at The University of Queensland.

## Author Contributions

AB performed the simulations, completed simulation analysis and helped write the paper.

CDC helped design the project, performed the electrophysiology experiments, generated Piezo1-APEX construct and helped write the paper.

JB assisted in simulation setup, wrote programs that were used in analysis, and contributed to the interpretation of the simulations.

JR, MB and RP carried out electron microscopy.

JL performed part of the electrophysiology experiments and generated Piezo1 mutations using site-directed mutagenesis.

JC performed qPCR and RNA isolation for microarray HSMC simulated and analysed the R808Q mutation.

MPH carried out lipidomics experiments and analysis. SJM helped design and interpret the simulations.

BM provided guidance for the project and helped interpret the data.

BC helped design the project and supervised the simulations, and assisted in the analysis and helped write the paper.

## Methods

### Molecular Dynamics Simulations

#### Structural Modelling

The structure of mPiezo1 (PDB ID 6B3R(Guo and MacKinnon, 2017)) was downloaded from the Protein Data Bank (https://www.rcsb.org). As some of the smaller loops were not resolved, these were built in using the program MODELLER (Fiser and Sali, 2003). Any large unstructured loops were not modelled. In addition, the sequence of the unresolved loops of mPiezo1 was run through PsiPred (http://bioinf.cs.ucl.ac.uk/psipred/). This determined that there is likely another amphipathic region before helix 33, in addition to the structurally resolved amphipathic helix. This helix was modelled into both structures using MODELLER(Fiser and Sali, 2003). The ‘full-length’ model starts from L577 and omits the following large extracellular loops: E718-D781, R1366-S1492, S1579-I1656, A1808-V1904. The model lacking the propeller domains starts from M1905 in the mouse model. For all simulation figures, the mouse sequence numbering has been converted to human numbering for ease of comparison to experimental data on human Piezo1.

#### Simulation Setup

GROMACS2018.1(Abraham et al., 2015) was used for the setup and execution simulations, with the martini2.2 forcefield (de Jong et al., 2013; Marrink et al., 2007) being used for all simulations. All simulations were run at 310K, with a van der Waals radius of 1.1nm and a timestep of 20fs (de Jong et al., 2016). The NVT simulations were run with isotropic pressure coupling. The NPT simulations used a Berendsen(Berendsen et al., 1984) thermostat, with semi-isotropic pressure coupling and pressure set to 1atm. Particle Mesh Ewald (PME) (Essmann et al., 1995) was used to calculate electrostatics.

Simulations were set up using the MARTINATE script, available on Github (https://github.com/Tsjerk/gromit). The initial homology model was coarse-grained using martinize (http://cgmartini.nl/index.php/tools2/proteins-and-bilayers) with an elastic network cutoff of 0.6nm. The elastic network was initially built using ElNeDyn(Periole et al., 2009), and extraneous elastic network bonds were removed using domELNEDIN(Siuda and Thøgersen, 2013). The protein was then embedded into a realistic mammalian membrane model using INSANE(Wassenaar et al., 2015) (see Supplementary Table 1 for the exact lipid composition), solvated with coarse-grain waters, neutralised and ionized with 0.15M NaCl. The size of the Piezo1-no propellers system is 50nm x 50nm x 25nm, which totals ∼500,000 coarse-grain particles. The size of the ‘full-length’ system is 60nm x 60nm x 25nm, which totals ∼ 710,000 coarse-grain particles. Both systems were then energy minimized using the steepest descent method for 5000 steps. After this initial minimization, position restrained – NVT was run for 1ps to remove possible high-energy interactions. Following this, three position-restrained – NPT equilibration steps were run for 5000 steps each, with 5fs, 10fs and 20fs timesteps respectively. Next, two initial NPT simulations were run for 1ps each, with a 20fs timestep. Finally, production runs for each system were run for 30μs in total. To allow the systems to equilibrate, all analysis was done using the last 10μs of the ‘full-length’ system and the last 15μs of the system lacking propeller domains.

For the mutation simulations, specifically R808Q and K2097E, the last frame of the first replicate of the ‘full-length’ wild-type simulation was manually mutated. As the mutations R808Q and K2097E involve removing one coarse-grain side chain bead, we manually deleted this bead and changed the name of the side chain. We then re-neutralised this structure and simulated each system for 10μs. For the Δ4K simulation, a new model was created using the ‘full-length’ structure as a template, but was built without K2166-K2169. This was simulated for 30μs, with the last 10μs of the simulation used for subsequent analysis.

Simulations under lateral tension were conducted by taking the last frame from each of the 30μs replicates, and applying tension using this starting point. With the exception of pressure, all inputs for these simulations are the same as above. In each of three repeat simulations, the lateral tension was increased in steps of −10 mMHg, with each step lasting 5 μs. Results shown represent the average height of each leaflet of the membrane determined from the last 3μs of all 3 replicates.

#### Simulation Analysis

Radial distribution functions were analysed using gmx rdf in GROMACS2018.1(Abraham et al., 2015). Curvature analysis and contact maps were calculated using in-house scripts. Depletion/Enrichment Indices were calculated using an updated version of the script used in (Corradi et al., 2018), and is available on the Martini website (cgmartini.nl). Thickness and density maps were calculated using modified versions of g_density and g_thickness(Castillo et al., 2013) (available for download from http://perso.ibcp.fr/luca.monticelli/tools/index.html), and compiled using GROMACS.4.5.6(Pronk et al., 2013). Visualisation was done using Visual Molecule Dynamics (VMD)(Humphrey, W., Dalke, A., Schulten, 1996).

#### Electrophysiology

Transiently transfected P1 KO HEK293T cells were plated on 35 mm dishes for patch-clamp analysis. The extracellular solution for cell-attached patches contained high K+ to zero the membrane potential and consisted 90 mM potassium aspartate, 50 mM KCl, 1 mM MgCl_2_ and 10 mM HEPES (pH 7.2) adjusted using KOH. The pipette solution contained either 145 mM CsCl or 145 mM NaCl with 10 mM HEPES (pH 7.2) adjusted using the respective hydroxide. EGTA was added to control levels of free pipette (extracellular) Ca^2+^ using an available online EGTA calculator—Ca-EGTA Calculator TS v1.3—Maxchelator. Negative pressure was applied to patch pipettes using a High Speed Pressure Clamp-1 (ALA Scientific Instruments) and recorded in millimeters of mercury (mmHg) using a piezoelectric pressure transducer (WPI, Sarasota, FL, USA). Borosilicate glass pipettes (Sigma, St Louis, MO, USA) were pulled using a vertical pipette puller (PP-83, Narashige, Japan) to produce electrodes with a resistance of 1.8-2.2 MΩ. Single-channel PIEZO1 currents were amplified using an AxoPatch 200B amplifier (Axon Instruments), and data were acquired at a sampling rate of 10 kHz with 1 kHz filtration and analyzed using pCLAMP10 software (Axon Instruments). Boltzmann distribution functions describe dependence of mesoscopic PIEZO1 channel currents and open probability, respectively, on the negative pressure applied to patch pipettes. The Boltzmann plots were obtained by fitting open probability *P*_o_∼*I*/*I*_max_ versus negative pressure using the expression *P*_o_/(1–*P*_o_)=exp [*α*(*P*–*P*_1/2_)], where *P* is the negative pressure (suction) [mmHg], *P*_1/2_ is the negative pressure at which *P*_o_=0.5, and *α* [mmHg^−1^] is the slope of the plot ln [*P*_o_/(1–*P*_o_)=[*α*(*P*–*P*_1/2_)] reflecting the channel mechanosensitivity.

#### Ca^2+^ imaging

Piezo1-1591-APEX2 expressing cells were loaded with Fura-2 (2 µM) in HEPES buffered Tyrodes solution with 1.5 mM CaCl_2_ for 30 minutes at room temperature. Imaging was carried out using a Nikon Tie2 microscope using a 20x objective illuminated with a COOLED p340 light source and analyzed using the advanced version of NIS elements (Nikon).

#### Microarray and qPCR

Total RNA was isolated from HEK293T cell lines using a standard Chloroform/Propanol/Ethanol precipitation protocol. Clariom D arrays were processed at the Ramaciotti Centre, University of New South Wales. The differentially expressed genes were determined using Affymetrix Transcriptome Analysis Console (TAC), following the software guidelines. T-test, Multiple Testing Corrections and False Discovery Rate Prediction were performed. The genes which showed at least a 1.5-fold difference between KO and one over-expressing cell line, with a p value of less than 0.05, were considered statistically significant and differentially expressed. Selected transcriptional differences were confirmed using qPCR. For qPCR one microgram of total RNA from each sample was reverse transcribed with the Super Script III First Strand Synthesis Super Mix Kit (Invitrogen, Carlsbad, California). The relative abundance of selected cDNA fragments was determined using a SYBR-Green Master Mix (Roche, Basel, Switzerland). Accumulation of polymerase chain reaction products and the threshold cycle were determined using the Bio Rad CFX 384 Real Time System, C1000 Touch Thermal Cycler. The primers used were as follows:

FAAH2_F_human: TGTGGTGGCATTACTGAAGG

FAAH2_R_human: ACAGCCAAGGGAAACTGAC

NAAA1_F_human: TACGACTTGGACTTGGTGCG

NAAA1_R_human: GACTCGTAGGCCAGGTTGAC

HPRT_F_human: TGAGGATTTGGAAAGGGTGT

HPRT_R_human: TAATCCAGCAGGTCAGCAAA

#### Cell culture

hP1 KO HEK293T cells were a kind gift from Ardem Patapoutian. The Piezo1-1591-GFP (referred to as over-expressor 1) and Piezo1-1591-mCherry HEK293T (referred to as over-expressor 2) were a kind gift from Philip Gottlieb (Maneshi et al., 2018).

#### Cloning

The human Piezo1 construct fused with GFP (Cox et al., 2016) was used to create the APEX2 fusion. In this construct the GFP inserted at amino acid position 1591 is flanked in the DNA sequence by a 5’ AgeI site and a 3’ SpeI site. A gene fragment of APEX2 (IDT technologies) was used with a native AgeI site ablated also flanked by a 5’ AgeI site and a 3’ SpeI site, digested and directly ligated into the Piezo1 SVLT vector. This Piezo1-1591-APEX2 construct was subcloned into pIRES2 for further use.

#### Electron microscopy

Piezo1 localization was visualized using either Piezo1-GFP or mCherry fusion constructs or Piezo1-APEX2 expressed in HEK293T Piezo1 KO cells or Cav1-KO mouse embryonic fibroblasts. For GFP and mCherry constructs it was necessary to co-transfect with GFP or mCherry-APEX binding proteins respectively. This was carried out using lipofectamine 3000. Localization of the APEX tag is fully and exhaustively documented elsewhere (Hall et al., 2018). Briefly, cells were fixed in 2.5% glutaraldehyde in 0.1 M cacodylate buffer for 30 min at room temperature and the 3,3′-diaminobenzidine tetrahydrochloride (DAB; 10 mg tablets; Sigma–Aldrich) reaction was carried out immediately after this in the presence of 5.88 mM H_2_O_2_ (Sigma–Aldrich). After fixation, cells were washed three times for 5 min in 0.1 M sodium cacodylate buffer. The DAB precipitate was contrasted by post-fixation with 1% osmium tetroxide (OsO4) for 2 minutes.

#### Cell lipid extraction

LC-MS grade solvents (water, chloroform, methanol, isopropanol), ammonium formate, ammonium acetate, butylated hydroxytoluene and formic acid (LC-MS grade) were purchased from Merck (Castle Hill, Australia). SPLASH® LipidoMix® internal standard mix was purchased from Avanti Polar Lipids (Alabaster, USA). Disposable 15 mL Pyrex® glass centrifuge tubes and Corning® phenolic caps with PTFE liners were purchased from Merck (Castle Hill, Australia).

Washed cell pellets from 5 HEK293T Piezo1 KO and 5 Piezo1-1591-mCherry overexpressing HEK2393T cultures were extracted using a chloroform:methanol:water-based modified Bligh-Dyer extraction procedure (Bligh and Dyer, 1959). Cell pellets were individually resuspended in ice cold 1.333 ml extraction methanol, comprising 14 ml of ice cold LC-MS grade methanol plus 0.01 mg/ml butylated hydroxytoluene as an antioxidant and 110 μl of SPLASH® LipidoMix® (i.e. ∼10 μl per sample). Resuspended pellets were drawn up using a clean single use glass Pasteur pipette and transferred to a clean disposable glass centrifuge tube. A volume of 0.666 ml of ice cold LC-MS grade chloroform was added to each tube and the mixture vortex-mixed for 30 seconds. The mixture was left to stand at 4°C for 60 minutes, after which it was vortex-mixed again for 10 seconds and a further volume of ice cold 0.666 ml LC-MS grade chloroform was added to each tube followed by 1.2 ml of ice cold LC-MS grade water. The mixture was then vortex-mixed for a further 30 seconds before centrifugation at 2900 r.c.f. for 20 minutes at 4°C. The lower (organic) layer was carefully retrieved using a clean single use glass Pasteur pipette and transferred to a clean 1.5 ml Eppendorf vial. A volume of 1 ml from each sample was split to two Eppendorf vials of 500 µl sample extract each for drying down of the organic sample extract using a Concentrator plus vacuum centrifuge (Eppendorf, Macquarie Park, Australia), employing the V-HV program with no heating (room temperature 22±1°C. The first of the dried split samples was resuspended in 200 µL of methanol/toluene (9:1 *v/v*) and this was vortex mixed for 10 seconds to dissolve/reconstitute the dried extract. This was then removed and transferred to the second tube and again this was vortex-mixed for 10 seconds to dissolve/reconstitute the second dried extract. Extraction blank samples were also prepared to determine solvent and reagent background. Quality control samples were prepared by pooling aliquots of each sample following the recommendation by Sangster et al. (2006)(Sangster et al., 2006). Final sample extracts, QCs and blanks were dispensed to HPLC vials with glass inserts. The sample lipidomes were then analysed by ultra-high performance liquid chromatography-quadrupole time-of-flight mass spectrometry.

#### Ultra-high performance liquid chromatography-quadrupole time-of-flight mass spectrometry (UHPLC-Q-ToF-MS)

Lipidomics analysis was performed using an Agilent 1290 Infinity II UHPLC system (Agilent Technologies, Santa Clara, USA) connected to a SCIEX TripleTOF 6600 quadrupole time-of-flight mass spectrometer equipped with a DuoSpray^TM^ ion source (SCIEX, Mulgrave, Australia). Chromatographic separation of lipid classes in the sample organic extracts was achieved using an Acquity® CSH C18 1.7 µm 2.1 x 100 mm column fitted with an Acquity® CSH C18 1.7 µm pre-column guard (Waters, Rydalmere, Australia). The mobile phases employed were A – 60:40 acetonitrile:water (*v/v*) and B – 90:10 isopropanol:acetonitrile (*v/v*) with the addition of different modifiers dependent upon the ionisation polarity (Cajka and Fiehn, 2016); in positive ionisation mode 10 mM ammonium formate and 0.1% formic acid was added to both mobile phases, in negative ionisation mode 10 mM ammonium acetate was added to both mobile phases. The chromatographic gradient employed for all experiments was as follows:

| Time (minutes) | %B |
| --- | --- |
| 0 | 15 |
| 3 | 30 |
| 4 | 60 |
| 12 | 80 |
| 16 | 99 |
| 17 | 99 |
| 17.1 | 2 |
| 18 | 2 |
| 19 | 99 |
| 19.5 | 99 |
| 19.7 | 15 |
| 24 | 15 |

The mobile phase flow rate was 0.5 ml/min, the column temperature was maintained at 60°C, the autosampler temperature was maintained at 4 °C and an injection volume of 2 µL was used for analysis in positive ionisation mode and 5 µL was used for analysis in negative ionisation mode. The mass spectrometer was operated using information dependent acquisition (IDA) in either positive or negative electrospray ionisation mode using the following parameters: curtain gas = 25; ion source gas 1 = 50 psi; ion source gas 2 = 60 psi; temperature = 500 °C; ion spray voltage floating = 5.5 kV (positive ion mode) and −4.5 kV (negative ion mode); declustering potential = (-)80 V; TOF MS accumulation time = 200 ms; TOF MS mass range = 200-1500 m/z; MS/MS accumulation time = 100 ms; collision energy = (-)35 eV; collision energy spread = (-)15 eV. The switch criteria for IDA were for ions between 200-1500 m/z exceeding 100 cps: maximum of 5 candidate ions to monitor per cycle, exclude former target ions for 5 seconds after 1 repeat, using dynamic background subtract. In addition a group-related pooled sample (i.e. KO and OE) was prepared and analysed using Sequential Windowed Acquisition of All Theoretical Fragment Ion Mass Spectra (SWATH-MS)(Gillet et al., 2012), a data-independent acquisition (DIA) method, using the following parameters: spectra were acquired with the LC-MS method as described above for IDA acquisition, using 34 mass isolation windows of width 30 Da, mass range of 200-1220 m/z, window overlap 1 Da; total cycle time was 3.36 seconds across a 24 minute run time (428 cycles). Instrument mass calibration was performed prior to the acquisition of each analytical batch and also automatically performed every 10 injections using an APCI positive/negative calibration solution via an external calibrant delivery system. The instrument was controlled using Analyst TF 1.7.1 (SCIEX, Mulgrave, Australia) and Analyst Device Driver via Manual/AAO Sync (Agilent Technologies, Santa Clara, USA).

#### Lipidomic data processing and analysis

As part of workflow development the data were processed using a number of different proprietary and open source software tools. Data were processed using MarkerView software v1.3.0.1 (SCIEX), MS-DIAL(Tsugawa et al., 2015), XCMS^plus^ (Smith et al., 2006; Tautenhahn et al., 2012) operating in a virtual machine environment and using default parameters for the SCIEX 5600+, and LipidMatch(Koelmel et al., 2017). For MS-DIAL processing, the proprietary SCIEX .wiff files were converted to analysis base files (.abf) using the Reifycs Abf Converter prior to processing. For XCMS^plus^ processing the .wiff and .wiffscan files were converted using the XCMS 3.6.3 converter within the VMWare virtual machine. LipidMatch uses the output results file from XCMS^plus^ as well as .ms2 files, which were converted from .wiff MS level 2 using ProteoWizard MSConvertGUI Version 3.0.18199 (Chambers et al., 2012). Processed data were analysed using XCMS^plus^, Excel (Microsoft Corporation, USA), SIMCA v15.0.0.4783 (Sartorius Stedim Data Analytics AB, Umeå, Sweden) and R/RStudio. Lipid nomenclature follows the recommendations proposed by the LIPID MAPS consortium(Fahy et al., 2009). Identifications derived from LipidMatch, XCMS and MS-DIAL were combined using the “CombineIDs.R” script provided in the LipidMatch download. Univariate analysis was performed within XCMSplus and Excel, with FDR correction following the method of Benjamin and Hochberg (1995)(Benjamini and Hochberg, 1995). Multivariate analysis was performed using SIMCA, including initial principal components analysis (PCA) followed by orthogonal projection to latent structures-discriminant analysis (OPLS-DA).

A total number of 9060, 11315 and 3977 positive ionisation variables and 8764, 8866 and 2594 negative ionisation variables were extracted using MS-DIAL, XCMS and MarkerView respectively. After univariate analysis of internal standard-corrected data (log transformation and Student’s *t*-test) this resulted in 112, 100 and 116 positive ionisation variables and 96, 126 and 70 negative ionisation variables passing the B-H FDR correction (set at 5%). A number of variables per analysis were selected based upon their OPLS-DA VIP_pred_ scores and univariate significance. The identification matches for these variables were combined across LipidMatch, MS-DIAL and XCMS and were visually curated using only those variables with associated MS/MS spectral data from the IDA experiments, as well as visual inspection of the chromatographic peaks.

