## Supplemental figures for "Piezo1 Induces Local Curvature in a Mammalian Membrane and Forms Specific Protein-Lipid Interactions"

### **Supplementary Information**

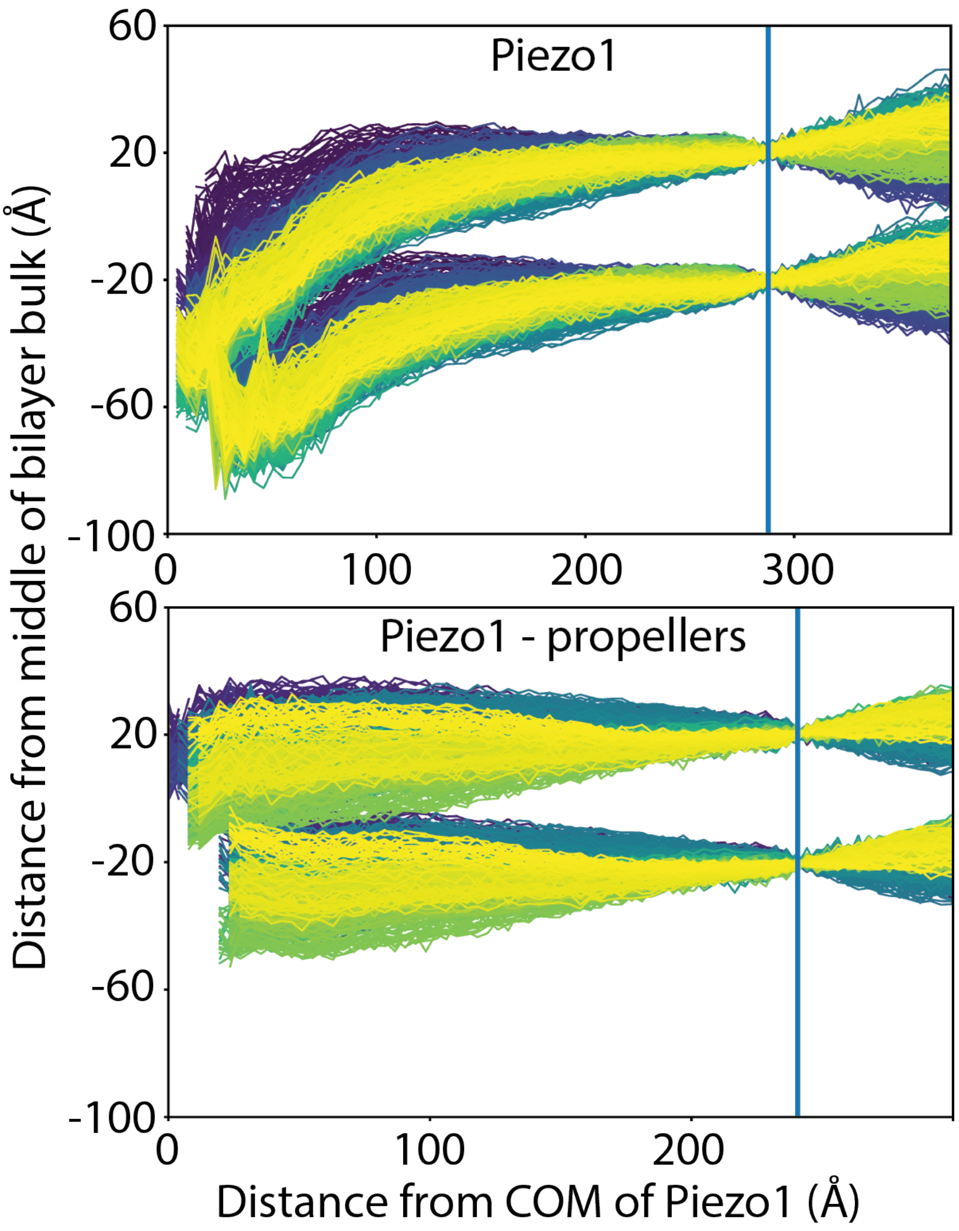


**Fig S1. Spontaneous curvature of the membrane occurs rapidly.** The distance from the middle of the bilayer bulk as a function of distance, shown for Piezo1 with (top) and without propellers (bottom) for the first microsecond of the 30µs simulation. Purple indicates the beginning of the simulation, whilst yellow indicates a frame close to 1µs of the simulation.

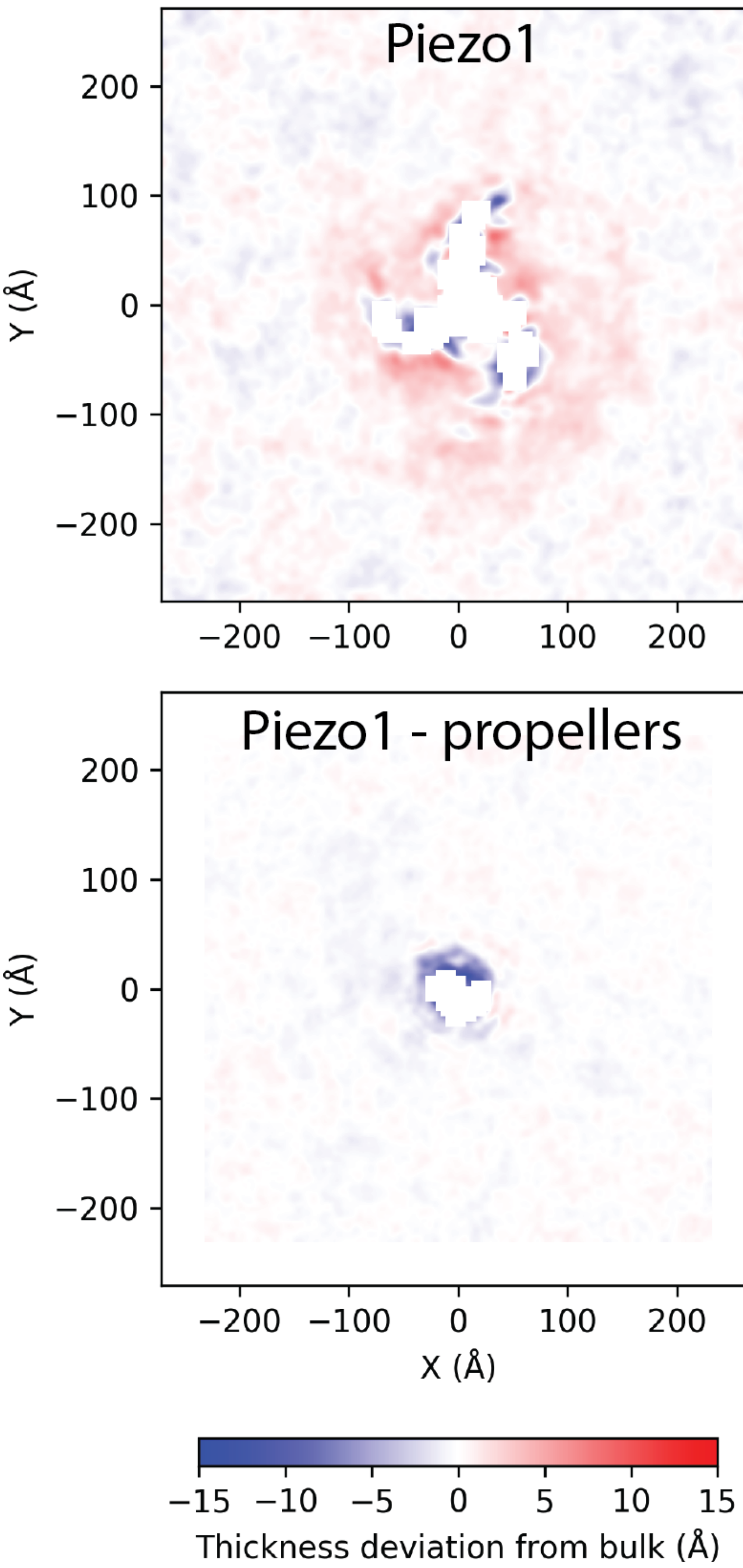


**Fig. S2. Piezo1's propellers increase the thickness of the surrounding bilayer.** The average thickness of the bilayer for Piezo1 with (top) and without (bottom) propellers.

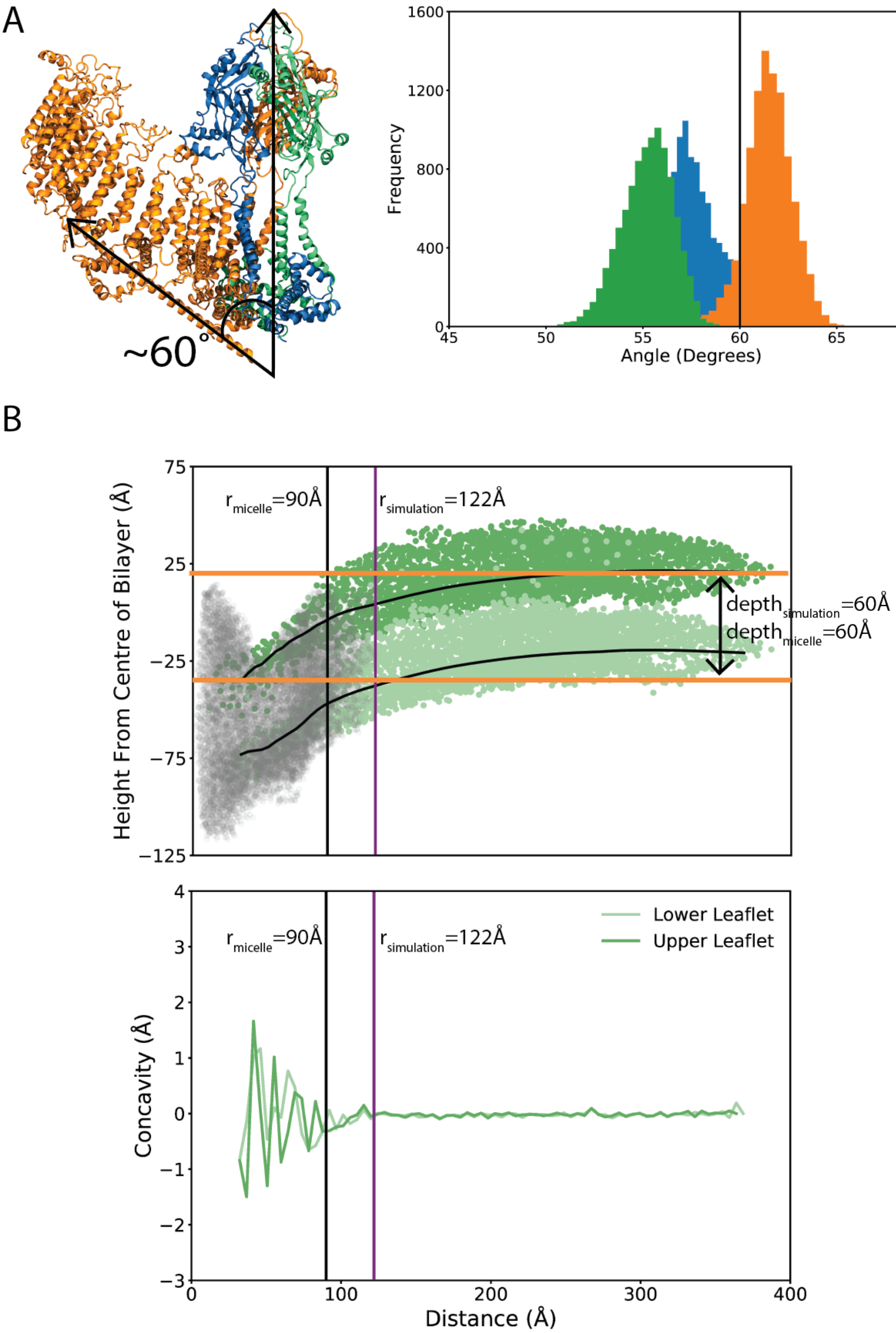


**Fig S3. Comparison between simulation and structure.** (A) The angle between the beam and the pore axis was shown to be ~60° in the structure from Mackinnon's group. (B) A histogram of the angle between the beam and the pore axis for all three monomers over the course of the last 10µs of simulation. (C) Average height of each membrane leaflet (as measured by the position of the phosphate atoms) as a function of distance from the centre of mass of the pore. Grey dots represent the average Cα atoms of Piezo1 and the dark and light green dots are the positions of the phosphate atoms of the top and lower leaflets of the last frame of each simulation system. The radius of curvature of the membrane deformation is indicated by the purple line, while the black line shows the equivalent measure from the 2017 structural paper of Guo and Mackinnon. The depth of the deformation is also indicated. (D) Quantification of the radius of curvature using the simulation system's concavity.

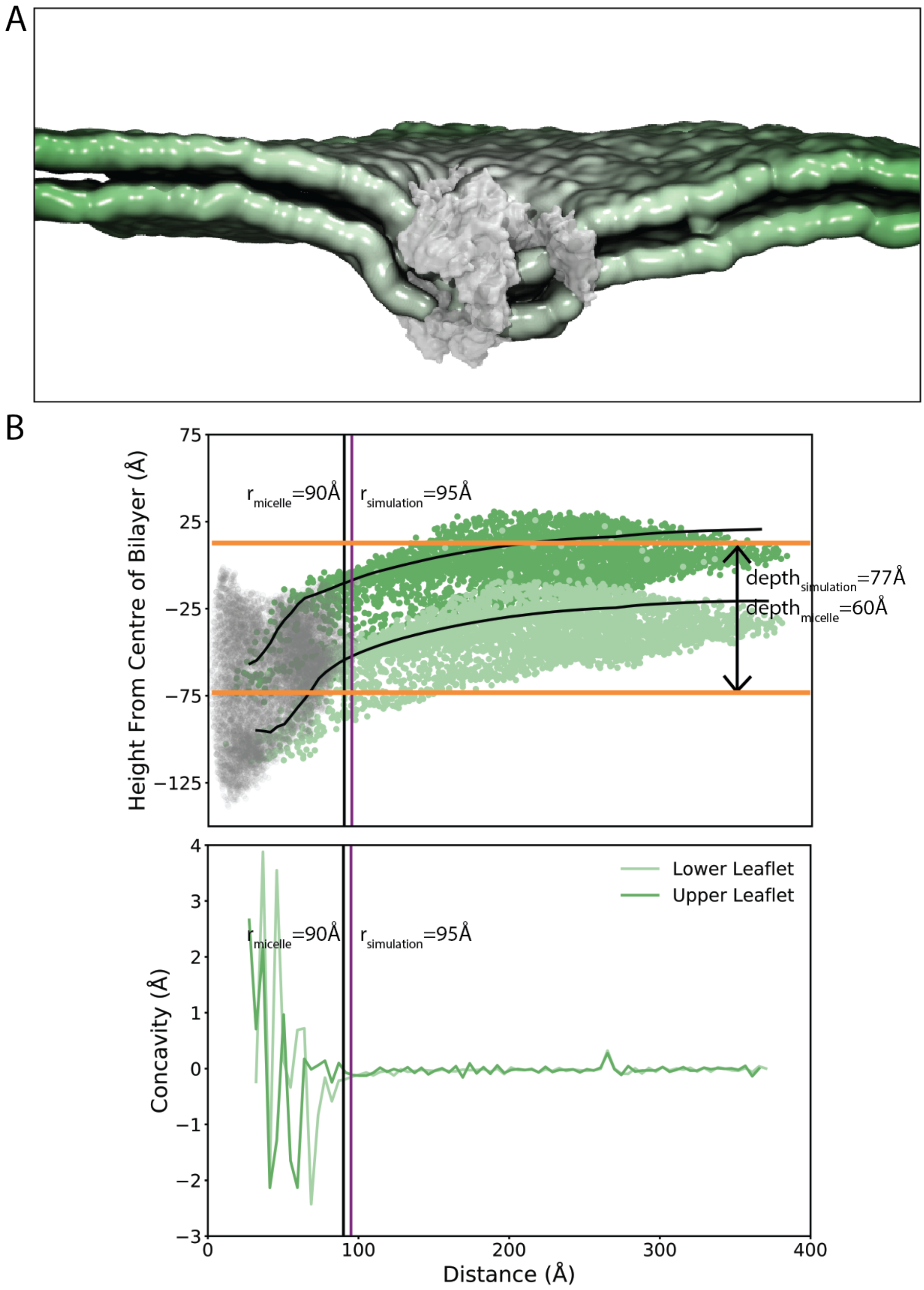


**Fig S4. Curvature in simulations lacking amphipathic helices.** (A) Representative image of the mouse Piezo1 simulation systems without amphipathic helices. The protein is represented by the grey surface, and the lipid phosphates are represented by green surfaces. Lipid tails, water and ions are omitted for clarity. (B) Average height of each membrane leaflet (as measured by the position of the phosphate atoms) as a function of distance from the centre of mass of the pore. Grey dots represent the average Cα atoms of Piezo1 and the dark and light green dots are the positions of the phosphate atoms of the top and lower leaflets of the last frame of each simulation system. The radius of curvature of the membrane deformation is indicated by the purple line, while the black line shows the equivalent measure from the 2017 structural paper of Guo and Mackinnon. The depth of the deformation is also indicated. (D) Quantification of the radius of curvature using the simulation system's concavity.

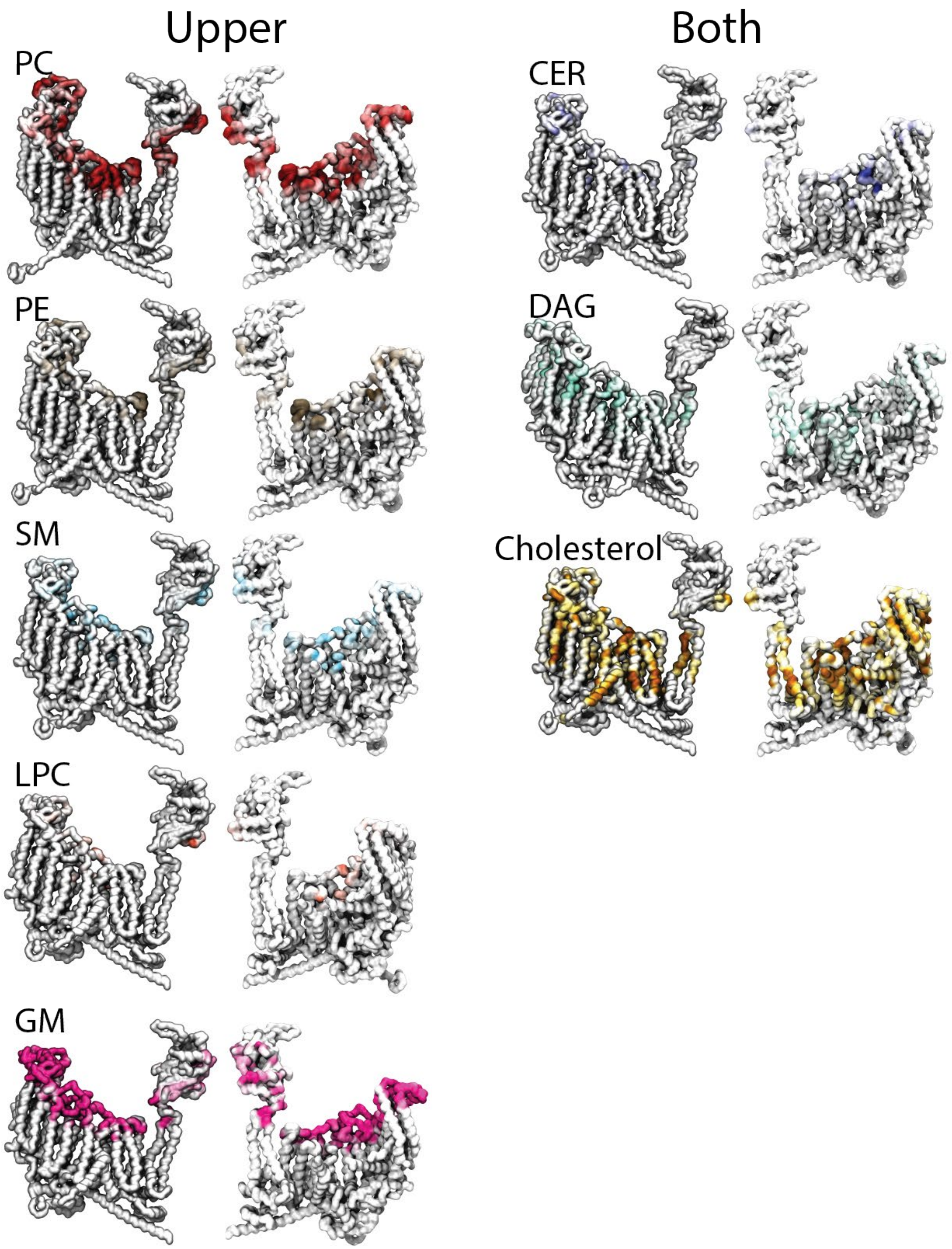


**Fig. S5. Significant contacts made with the lipids either in the upper leaflet or both leaflets.** Significant lipid contacts are mapped onto a monomer of Piezo1 and show the location and propensity of lipids to interact with Piezo1. This includes lipids in the upper leaflet (left) or lipids that flip-flop between both leaflets (right).

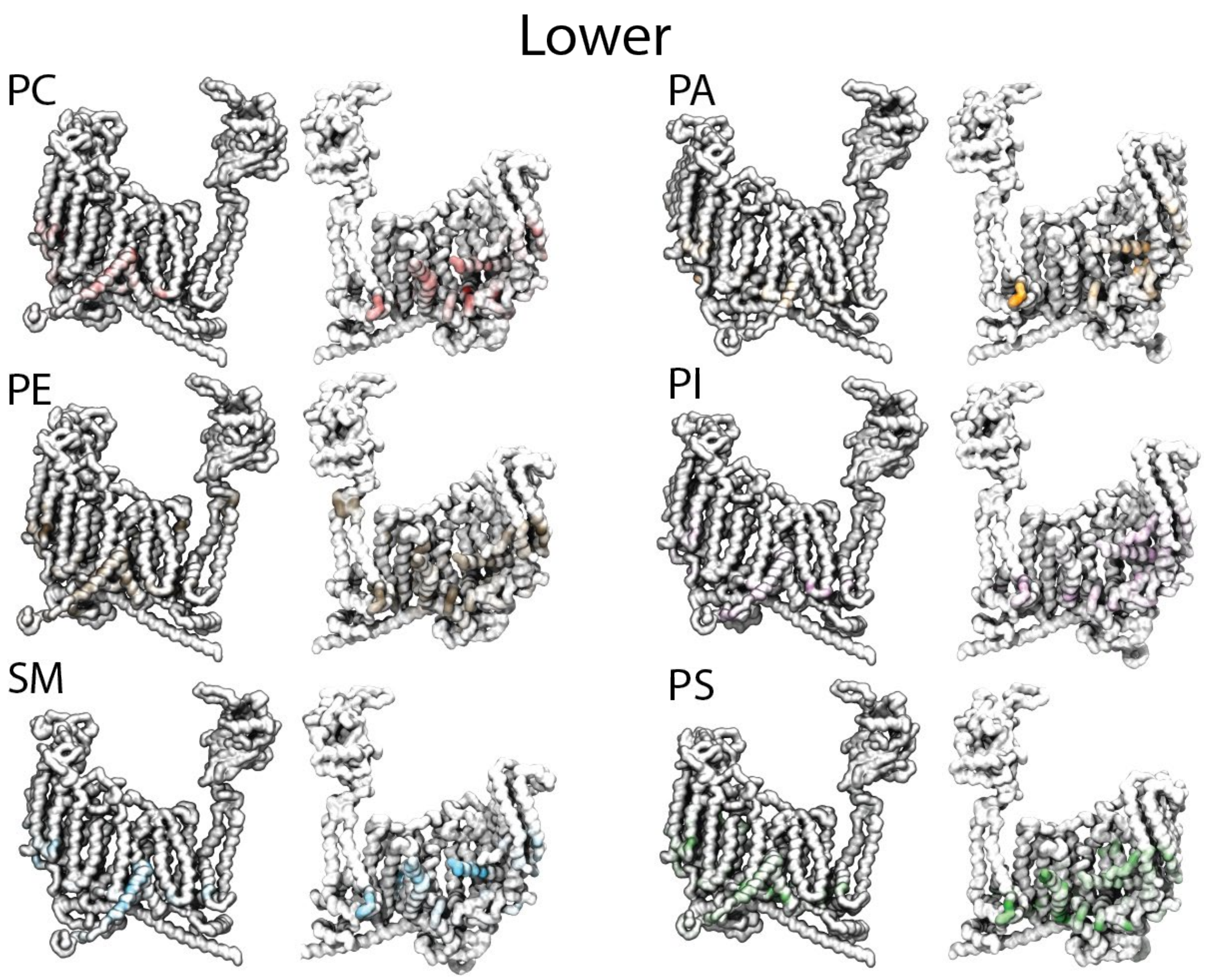


**Fig. S6. Significant contacts made with the lipids in the lower leaflet.** Significant lipid contacts are mapped onto a monomer of Piezo1, and show the location and propensity of lipids in the lower leaflet to interact with Piezo1.

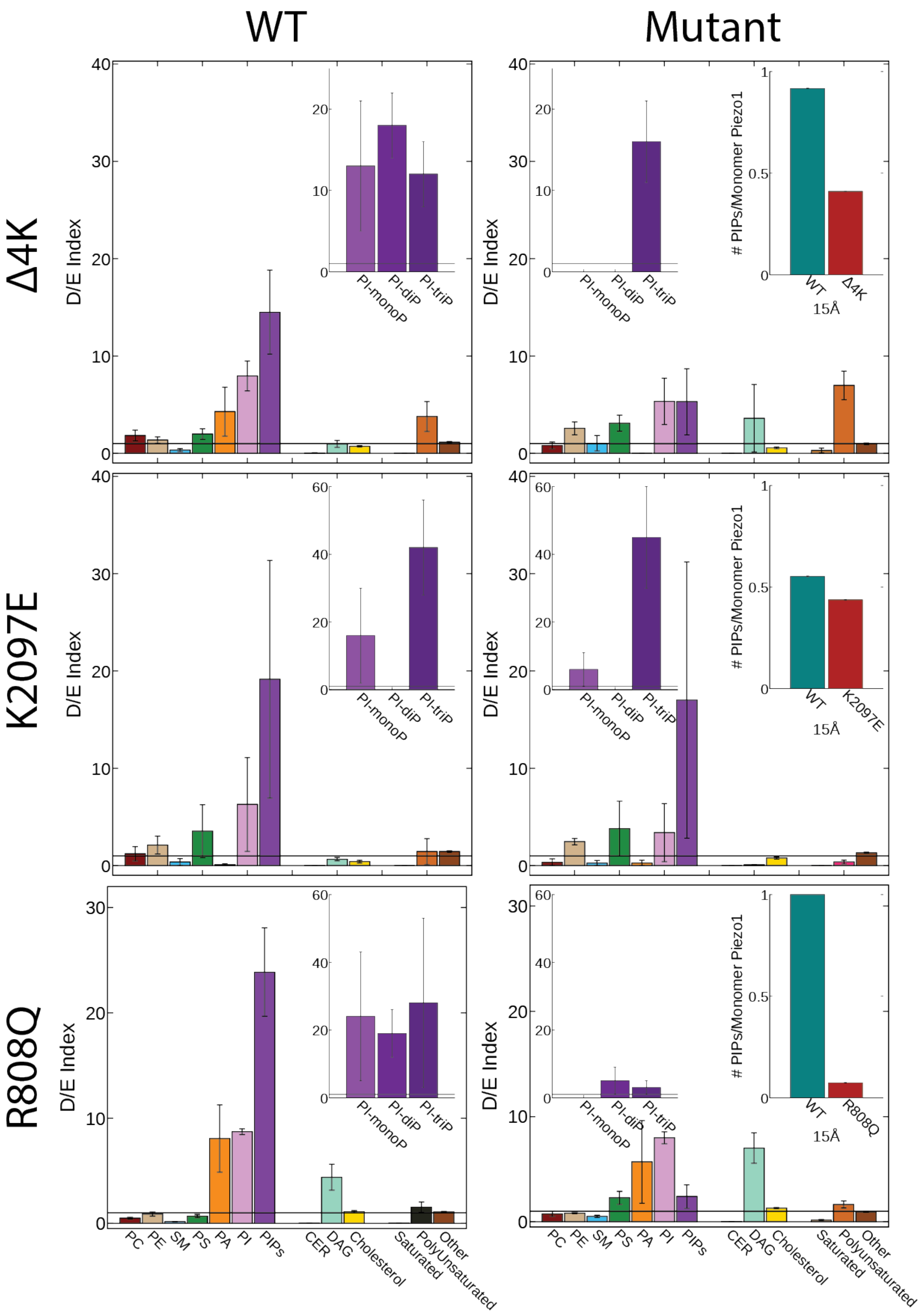


**Fig. S7. Mutation of disease causing residues alters PIP binding.** Depletion/Enrichment (D/E) Indices for both the WT (left) and mutants (right) for the Δ4K (top row), K2097E (middle row), and R808Q (bottom row) mutants. Averages of the Δ4K, middle and bottom rows of D/E indices were calculated by averaging each propeller's D/E index for each lipid, and the SEM is shown. The WT Δ4K mutation is an average of three replicates, which each had their own average and SEM from the three propellers in each simulation. The insets show D/E for different PIP species, and the number of PIP molecules per monomer within 15Å of the site of mutation in both the WT and mutant.

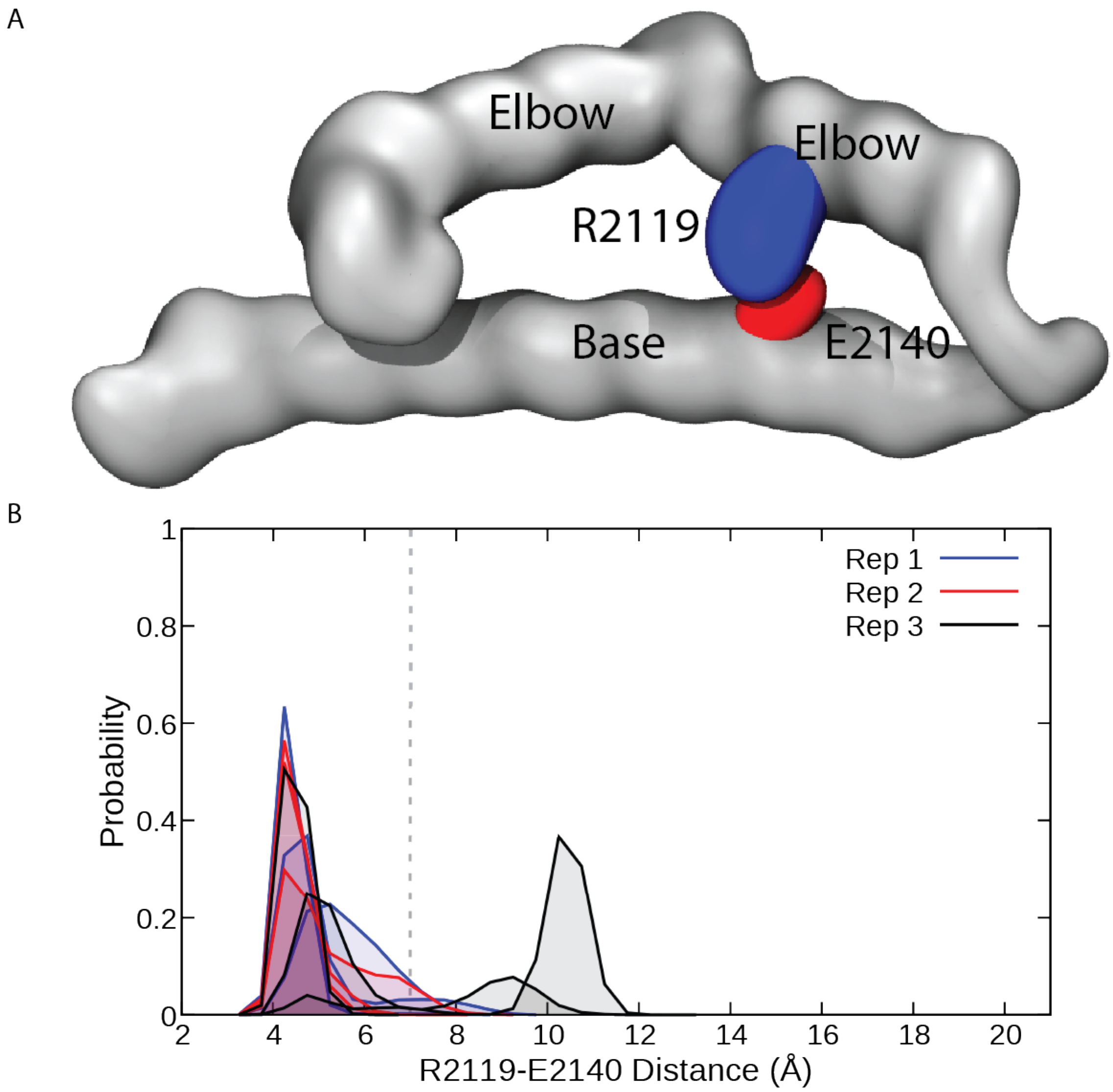


**Fig. S8. R2119 forms a salt bridge with E2140**. (A) Visualisation of R2119 forming a salt bridge with E2140. The backbone of the protein is in grey surface, with arginine and glutamate in blue and red, respectively. (B) Histogram of the distance between R2119 and E2140 for each monomer in each replicate for the final 10µs of the simulation.

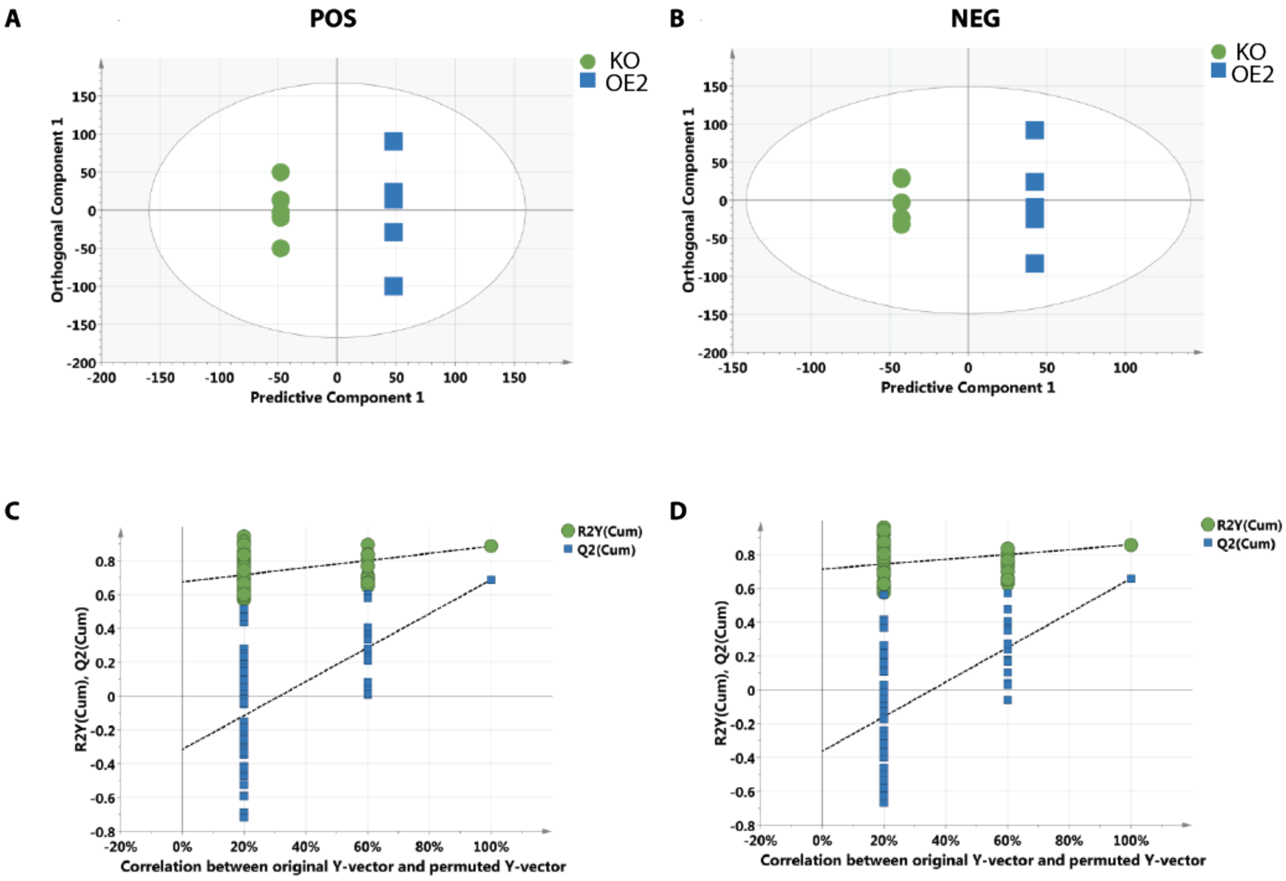


**Fig. S9. Multivariate analysis of the lipidome from the knockout and over-expressor cell lines**. (A) OPLS-DA scores plot of positive ionisation data from KO vs OE2 [Piezo1-1591-mCherry] (1+4 components; $R^2X[cum] = 0.224$; $R^2Y = 1$; $Q^2[cum] = 0.905$). (B) OPLS-DA scores plot of negative ionisation data from KO vs MC (1+4 components; $R^2[cum] = 0.225$; $R^2Y = 1$; $Q^2[cum] = 0.870$); C) Model validation permutation plot (100 permutations) for OPLS-DA of positive ionisation data from KO vs OE2 [Piezo1-1591-mCherry]. (D) model validation permutation plot (100 permutations) for OPLS-DA of negative ionisation data from KO vs OE2 [Piezo1-1591-mCherry]. All data were $log_{10}$-transformed prior to autofitted model generation.
